# “Non-viral CRISPR/Cas9 Mutagenesis for Streamlined Generation of Mouse Lung Cancer Models.”

**DOI:** 10.1101/2023.12.21.572771

**Authors:** Irene Lara-Saez, Angeles Mencia, Enrique Recuero, Yinghao Li, Marta García, Marta Oteo, Marta I Gallego, Ana Belen Enguita, Diana de Prado-Verdún, A Sigen, Wenxin Wang, Ramón García-Escudero, Rodolfo Murillas, Mirentxu Santos

## Abstract

Functional analysis in mouse models is necessary to establish the involvement of a set of genetic variations in tumor development. Many lung cancer models have been developed using genetic techniques to create gain- or loss-of-function alleles in genes involved in tumorigenesis; however, because of their labor- and time-intensive nature, these models are not suitable for quick and flexible hypothesis testing. Here we introduce a lung mutagenesis platform that utilizes CRISPR/Cas9 RNPs delivered via cationic polymers. This approach allows for the simultaneous inactivation of multiple genes. We validate the effectiveness of this system by targeting a group of tumor suppressor genes, specifically *Rb1*, *Rbl1*, *Pten*, and *Trp53*, which were chosen for their potential to cause lung tumors, namely Small Cell Lung Carcinoma (SCLC). This polymer-based delivery platform enables the modeling of lung tumorigenesis independently of the genetic background, thus simplifying and expediting the process without the need for modifying the mouse germline or creating custom viral vectors.

**Significance:** The development of models to rapidly introduce gene mutations into lung tissue to study their impact on tumor growth is critical for advancing the functional genomics of lung cancer. While previous methods using viral vectors and genetic manipulation in mice have been time-consuming and expensive, here we describe a new technique using cationic polymers as non-viral carriers for CRISPR/Cas9 delivery to induce cancer driving mutations that streamlines this process. This approach mimics natural mutations in lung cancer and accelerates the generation of accurate tumor models. Our study demonstrates the effectiveness of this method in generating small cell lung cancer (SCLC) by modifying four tumor suppressor genes in different mouse genetic backgrounds. This innovative strategy holds promise for faster and more cost-effective cancer modeling.

**Graphical Abstrac:** Small Cell Lung Cancer (SCLC) tumors are rapidly generated in any mouse genetic background by using cationic polymers to simultaneously deliver Cas9 and gRNAs targeting the *Rb1*, *Rbl1*, *Pten* and *Trp53* tumor-suppressor genes to the adult airway respiratory system *in vivo*. Addition of the frt guide in the RC::FLTG mice provides a tdTomato gene editing reporter. This study shows the feasibility of rapidly generating lung cancer mouse models via somatic genome engineering through delivery of all CRISPR components in the form of nanoparticles.

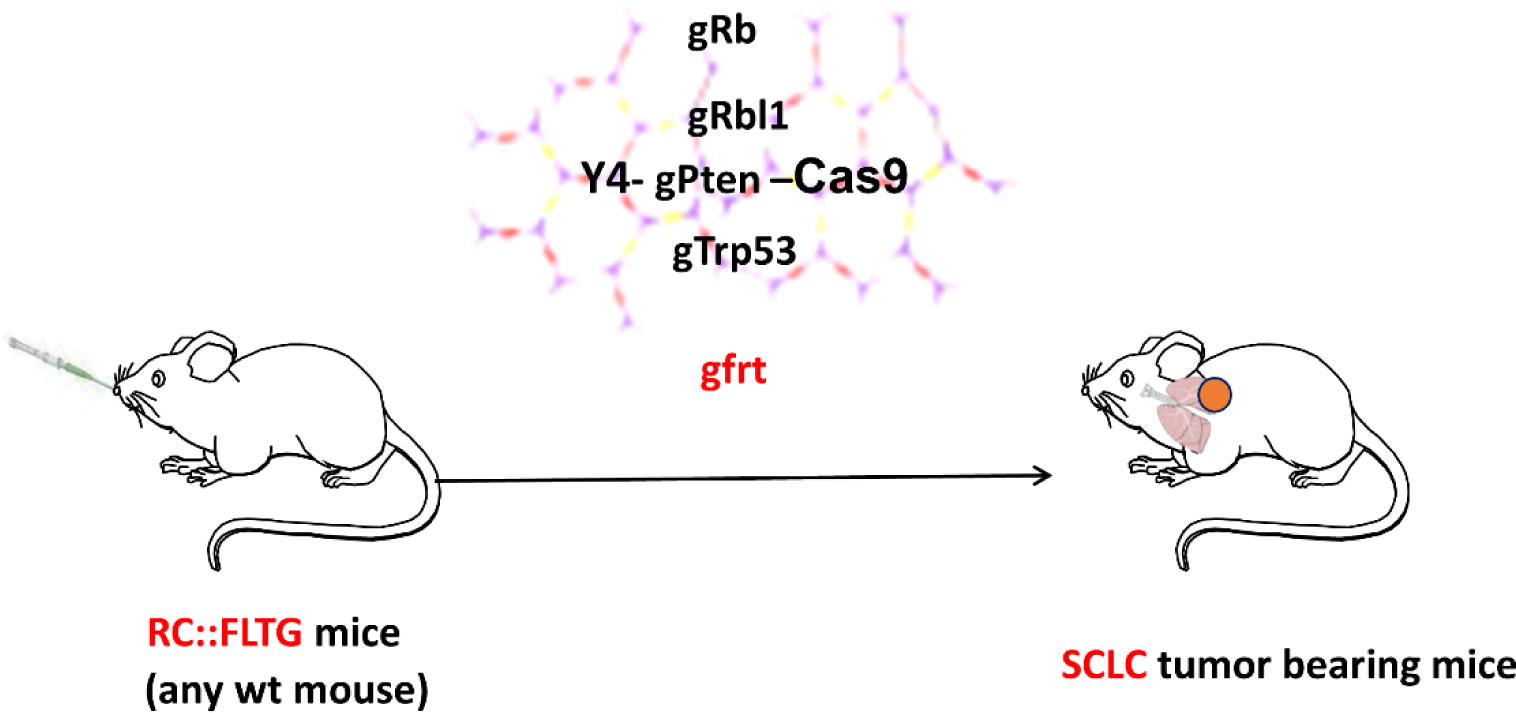

## INTRODUCTION

Large-scale genomic analysis of lung tumors has identified many recurrent gene mutations with unclear roles in tumor development ([1–3]. Therefore, it is essential to generate models that facilitate the rapid introduction of sets of mutations into lung tissue to assess their importance in disease progression. Over the years, the mouse modeling community has developed sophisticated tools for modeling cancer in mice, including transgenesis to achieve tissue-specific gene expression alterations and the introduction of mutations in target genes using mouse embryonic stem cells [4]. Precise gene inactivation in adult somatic tissues through germline alleles with sequences flanked by loxP can be achieved using adenoviral and lentiviral vectors expressing Cre recombinase. This method allows time-controlled, tissue-specific manipulation [5] and has provided important insights into the effect of gene inactivation on tumor development [6–9]. However, breeding mice to generate complex genotypes with different engineered alleles takes years, involves many animals and is extremely costly, which is a major drawback.

New nuclease-based gene editing techniques, such as CRISPR, facilitate the introduction of insertions and deletions at specific positions with high precision and efficiency, and allow the rapid creation of tumorigenesis models that faithfully recapitulate key features in the initiation and development of human cancer [10,11]. A combination of Cre-mediated oncogenic activation and CRISPR-mediated editing of tumor suppressor genes shed light on gene function in lung adenocarcinoma development in different studies in which the use of viral vectors and germline engineering was required [10,12,13]. Similar methods were used to model tumor suppressor gene (TSG) inactivation leading to Small Cell Lung Cancer (SCLC) [14] and an oncogenic chromosomal rearrangement causing non-Small Cell Lung Cancer (NSCLC) [15].

The direct delivery of the CRISPR system into tissues holds significant potential for establishing animal models through *in vivo* generation of somatic genetic modifications, thereby eliminating the requirement for additional genetic tools. The ability to directly change tumor suppressor genes and produce point mutations in oncogenes *in vivo* was proven by hydrodynamic delivery of a plasmid for sgRNAs and Cas9 expression into the liver [16]. An important breakthrough in gene editing was the utilization of CRISPR/Cas9 ribonucleoproteins (RNPs) delivered via electroporation, which enables highly efficient targeted modifications in genomes. The administration of CRISPR/Cas9 RNPs leads to immediate chromosomal DNA excision, followed by their rapid degradation within cells, thereby minimizing off-target effects. *In vitro* electroporation of RNPs is highly effective for gene editing of many different cell types [17,18]. *In vivo* electroporation of CRISPR/RNPs for inactivation of tumor suppressor genes in uterine tissue resulted in endometrial tumor formation [19] and lipid nanoparticles formulated with different ratios of permanently cationic and ionizable lipids proved useful to deliver CRISPR/Cas9 RNPs to the lungs after intravenous injection to produce oncogenic translocations causing NSCLC tumors [20]. The development of non-viral carriers that allow direct delivery of CRISPR/Cas9 RNPs into the lung may facilitate the establishment of cancer models for rapid functional genomic research [20].

Cationic polymers possess a number of advantageous characteristics for functioning as nucleic acid carriers, including biosafety, simple synthesis, substantial cargo-carrying capacity and biodegradability [21]. Polymeric nanoparticles are formed by electrostatic reactions when combining nucleic acids and aqueous solutions of cationic polymers. This protects nucleic acidsfrom degradation and facilitates their cellular uptakeby endocytosis. The escape from the endosomal compartment is due to the proton sponge effect, then polymers degrade hydrolytically in the cytoplasm releasingtheir cargo [22]. Lynn et al. first reported the synthesis of linear poly(β-aminoesters) (PBAEs) and highlighted their potential for gene delivery [23]. Later, a screen of a large library of linear PBAEs [24] identified a particularly efficient compound for gene transfection resulting from end-capping a C32 backbone with 1,3-diaminopropane (103) [25]. Other groups synthesized branched versions of poly(β-aminoesters) to explore the possibilities of improved complexation with nucleic acids [26]. In previous work, we have demonstrated the ability of these cationic polymers to deliver CRISPR reagents in both *in vivo* and *in vitro* gene editing applications [27–29]. Furthermore, this delivery system enables the simultaneous targeting of multiple genes using readily accessible and cost-effective commercial reagents, making direct CRISPR/Cas9 delivery to lung tissue a promising and straightforward method for introducing mutations in cancer-related genes. This underscores the value of polymeric vectors as an alternative approach for delivering CRISPR RNPs [29,30].

In this study, we describe a model of SCLC through the inactivation of four tumor suppressor genes by *in vivo* gene editing using CRISPR/Cas9 RNPs delivered by polymeric carriers. Furthermore, we show the efficacy of this rapid cancer modeling platform across various pure mouse genetic backgrounds.

## RESULTS

### *In vivo* delivery of CRISPR/Cas9 RNPs ribopolyplexes in the respiratory system

Our first goal was to establish a mouse model in which to test CRISPR/Cas9 RNPs-mediated gene editing *in vivo*. The RC::FLTG dual recombinase-response indicator allele contains an frt-flanked transcriptional STOP sequence that prevents transcription of tdTomato. *Flp* recombinase expression results in tdTomato fluorescence [31]. We designed gRNAs to target the frt sequence by selecting the 20 nucleotide sequence prior to the protospacer adjacent motif (PAM) on both the sense (guide 1) and antisense (guide 2) strands. These gRNAs were tested by electroporation of their CRISPR/Cas9 ribonucleoproteins into mouse embryonic fibroblasts (MEFs) derived from RC::FLTG mice for their ability to remove the transcriptional STOP sequence and induce tdTomato expression. Red fluorescence was detected in approximately 50% of the cells, demonstrating that this allele can be used as a reporter for gene editing with a gRNA targeting the frt sequence. Since guide 1 worked slightly better than guide 2 we selected guide 1 for further experiments (Fig1).

**Fig. 1.**
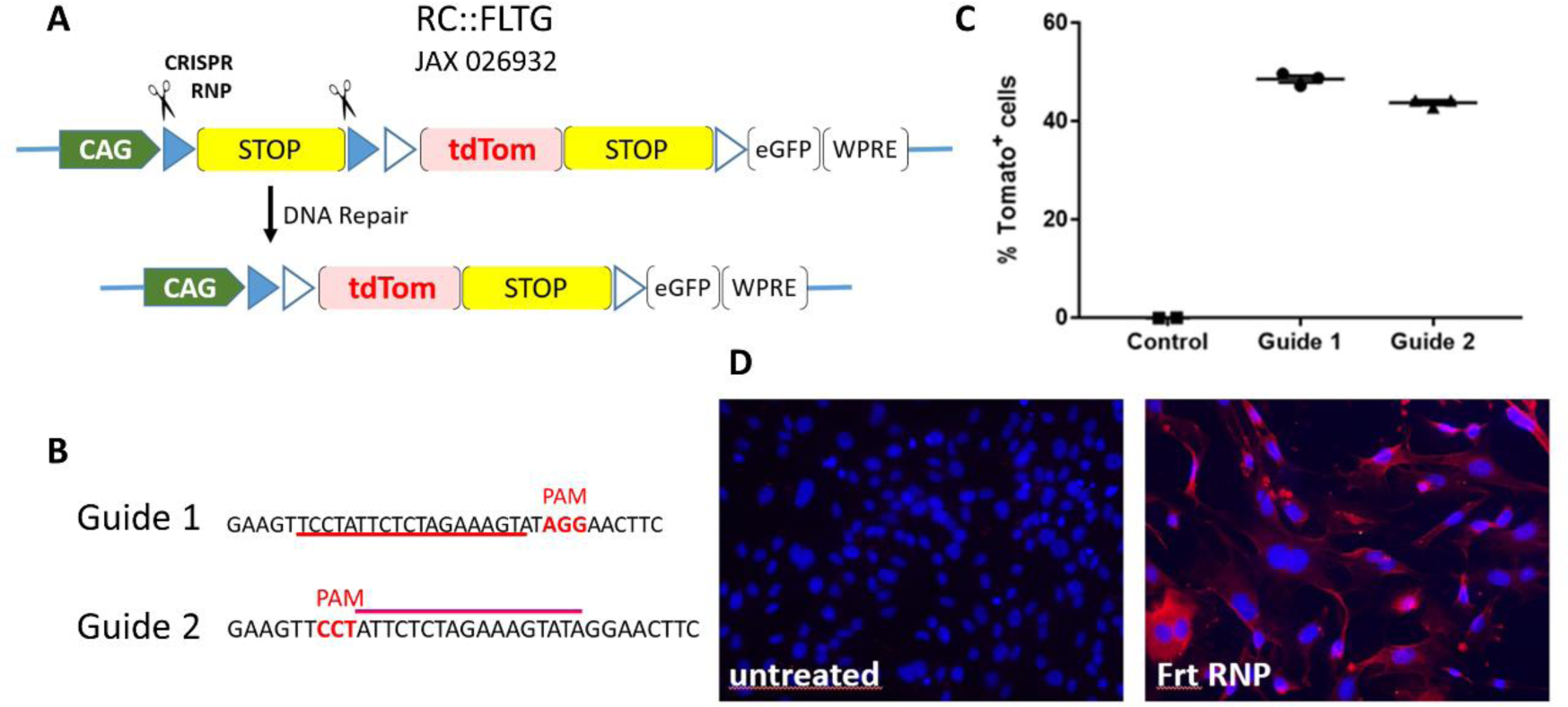
A system to assess *in vivo* gene editing in mouse models. A) The dual-recombinase-responsive fluorescent indicator allele, *RC::FLTG* (Rosa-CAG-frt-loxP-tdTomato-eGFP) (Jackson Lab, 026932). frt (flp recognition target sequence) is depicted as blue triangles. B) gRNAs on frt sequence. C) Flow cytometry analysis and (D) red fluorescence detection of tdTomato expression after nucleofection of mouse embryonic fibroblasts from RC::FLTG mice with CRISPR/Cas9 RNPs for frt guides.

To establish the feasibility of performing in vivo mutagenesis in the respiratory tract, we decided to evaluate a family of PBAE polymers for their capability to serve as carriers of CRISPR RNPs in the respiratory airways. Specifically, we tested Y4 Linear, which has been previously described [24,25] as C32-103, along with two novel derivatives not documented elsewhere: Y4 L5 (branched) and Y4 Cys (branched and modified with Cys) (see Fig. S1). Y4 L5 is a branched version of Y4 achieved by introducing a 4-branching diamine, while Y4 Cys is a branched version that incorporates cysteine residues for end capping, which provides both positive and negative charges to potentially enhance binding to the protein component of CRISPR/Cas9 RNPs. To monitor the synthesis reaction of the polymers and confirm their chemical structure, Nuclear Magnetic Resonance (NMR) and Gel Permeation Chromatography (GPC) techniques were used showing that the three polymers had comparable molecular weights and PDIs (Mw from 16 kDa to 18k Da, Polydispersity Index (PDI) from 4.32 to 5.4 (Fig. S2).

After assembly of ribopolyplexes consisting of the polymers to be evaluated mixed with CRISPR/Cas9 RNPs for frt gRNA, a single administration was performed in the respiratory tract of RC::FLTG mice by intratracheal intubation. Six to eight days after delivery, trachea and lungs were dissected and processed for immunohistochemistry to detect tdTomato. Tomato positive cells were detected along the respiratory epithelium, including the trachea, bronchi, bronchiole and parenchyma (Fig. 2 A-E). As a control, in the absence of polymer, no staining was observed in RC::FLTG mice tested with the frt guide and CRISPR components, or in wild type C57BL/6J mice tested with the ribopolyplexes (n=2 for each type of mouse and polymer tested)(Fig. 2 F,G). Out of a total of 27 mice that received Y4 polyplexes, we observed areas with tomato-positive cells in 7 mice: 6 out of 13 treated with the branched Y4 (Y4 L5), 1 out of 7 in the Y4 Linear-treated group, and none in the mice treated with the Cys modified version (0 out of 7).

**Fig. 2.**
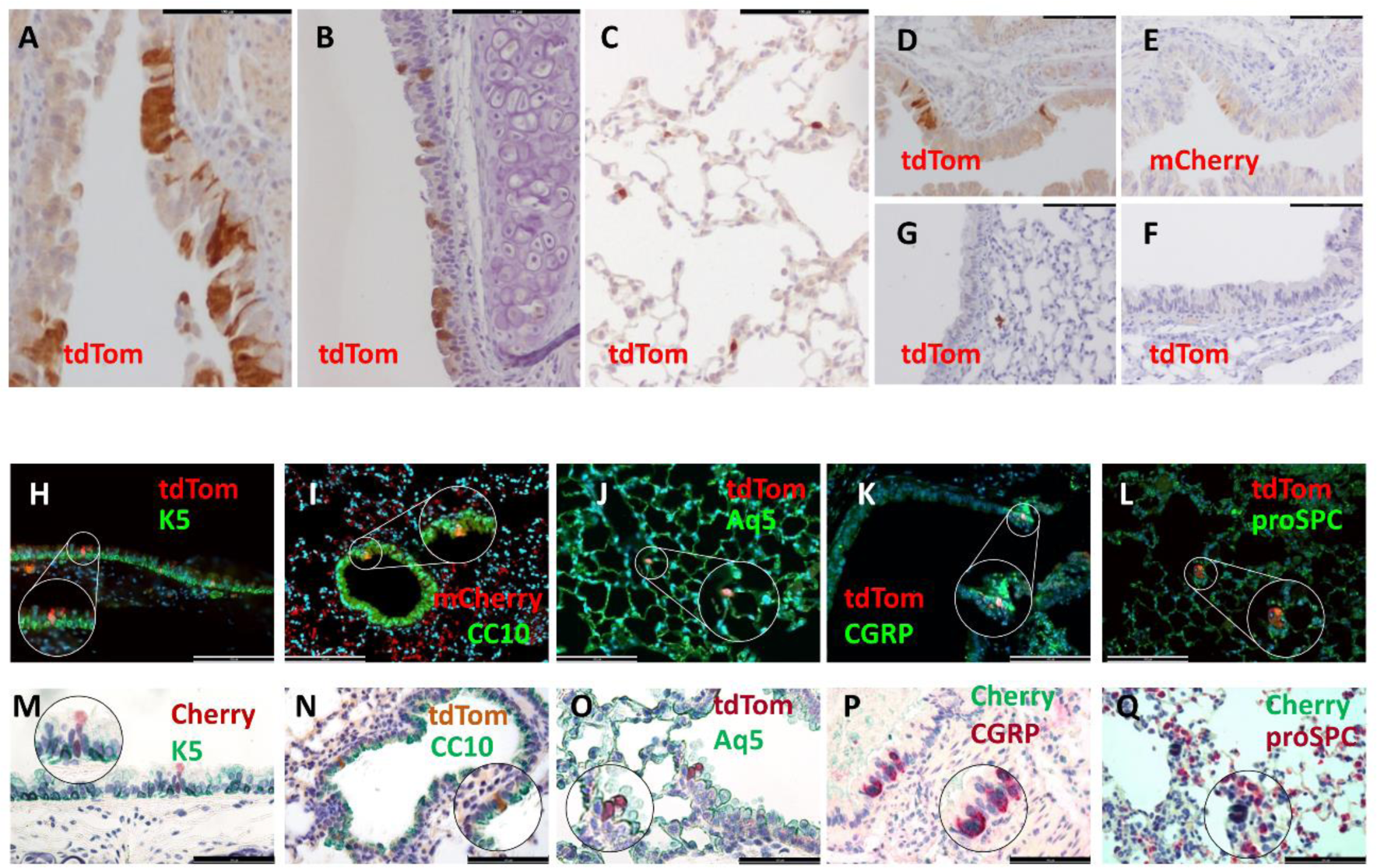
Efficient *in vivo* gene editing in the reporter mice airway system. *In vivo* gene editing in adult RC::FLTG lung cells assessed by immunohistochemical detection of tdTomato protein in bronchi (A), trachea (B) parenchyma (C) and bronchiole (D, E). IHQ with anti TdTomato (A-D, F, G) or anti mCherry (E) antibodies in paraffin-embedded respiratory tissue at week 1 after delivery of polyplexes. C57BL wt mice showed no expression of reporter protein (F, G) F: Trachea, G: bronchiole and parenchyma. Double immunofluorescence (H-L) and Dual color immunohistochemistry (M-Q) of the quoted proteins. BARS= 100µm.

To identify the different cell types targeted by this procedure, we performed a marker expression analysis of lung cell types. Double immunofluorescence staining revealed that the respiratory cells in which the ribopolyplex-mediated gene editing had occurred were keratin K5 expressing basal cells abundant in trachea and bronchi (Fig. 2H) as well as lung epithelial cells including Clara/club (Fig. 2I), alveolar type 1(Fig. 2J) neuroendocrine (Fig. 2K) and alveolar type 2 (Fig. 2L) cells. Further confirmation of the diversity of cell types targeted was achieved using dual color immunohistochemistry (Fig. 2M-Q).

The presence of the tomato reporter protein proved *in vivo* lung gene editing following polymer-mediated RNP delivery. The scattered distribution of edited cells was similar to that observed after delivery of Cre recombinase using adenoviral vectors [6]. The efficiency and diversity of targeted cell types observed in this gene editing pattern indicated that it may be possible to target a sizable population of cancer-initiating cells. This prompted us to explore the feasibility of using this non-viral gene editing system to generate lung cancer models.

### Lung carcinogenesis using CRISPR/Cas9-Y4 polyplexes for tumor suppressor gene inactivation

We have previously demonstrated that simultaneous inactivation of *Rb1, Rbl1, Pten* and *Trp53* genes by means of adenoviral *cre* infection in conditional *cre/loxP* mouse models leads to the development of high-grade neuroendocrine lung carcinomas [6,32,33]. We therefore designed guides to inactivate *Rb1*, *Rbl1* and *Pten* by gene editing with CRISPR/Cas9 ribopolyplexes and used a previously published guide to disrupt the *Trp53* gene [34]. The effectiveness of the guides for generating indel mutations was initially assessed by nucleofection of RC::FLTG MEFs (Fig. S3) with CRISPR/Cas9 RNPs either individually or as a set of four guides (crRNA for *Rb1*, *Rbl1*, *Pten* and *Trp53*, hereafter designated as Q). The specific region targeted for mutation in each gene was PCR amplified and analyzed through Sanger sequencing, revealing high frequencies of indels at the intended positions, as evidenced by tracking of indels by decomposition (TIDE) analysis. The spectrum of indels found for the nucleofections of the RNPs was the same for each guide individually or as a Q-set, indicating that the editing efficiency did not decrease in the multiplex experiment (Fig. S3).

Given that CRISPR/Cas9 RNPs effectively inactivate these tumor suppressor genes in RC::FLTG MEFs (Fig. S3), we administered the RNPs/polymers nanoparticles (i.e the ribopolyplexes) to the respiratory system of adult RC::FLTG mice by intratracheal intubation. We also added the frt guide to the complex as a marker of edited cells. Overall we treated 35 RC::FLTG mice with Y4 polyplexes: 15 with Y4 Linear, 10 with Y4 L5 and 5 with Y4 Cys polyplexes (Table 1). To assess and follow up on the development of tumors following inoculation, the mice underwent bimonthly monitoring using computed tomography (CT) imaging. Developing tumors could be detected as masses growing in the lungs or trachea (Fig. S4). When mice displayed respiratory distress or exhibited symptoms of disease, such as weight loss, lethargy, or hunched posture, they were euthanized and subjected to necropsy procedures. Mice from the *Qfrt Y4Linear* group (n=15) developed tumors with a frequency of 70% and a latency period of 4-12 months (Fig. 3A, B, Table S1). Tumors appeared at 5–6 months in mice from *Qfrt Y4L5* group (*n* = 10), with an incidence of 30%. One mouse (out of five) of the *Qfrt Y4Cys* group developed tumors 14 months after polyplex delivery (incidence 20%). Strong red fluorescence emission was observed in certain tumors that developed after delivery of ribopolyplexes, providing evidence of marker allele *in vivo* gene editing (Fig. 3C).This was further confirmed by immunohistochemistry (Fig. 3 D).

**Table 1.** Tumorigenesis output in experimental groups.

| Group | RNPs |  |  | polymer | receptor mice | tumors | # mice | #mice with tumors |
| --- | --- | --- | --- | --- | --- | --- | --- | --- |
|  | crRNA | tracr | *cas9* |  |  |  |  |  |
| Qfrt Y4Linear | Q frt | yes | yes | Y4 Linear | RC::FLTG | yes | 15 | 10/15 |
| Qfrt Y4L5 | Q frt | yes | yes | Y4 L5 | RC::FLTG | yes | 10 | 3/10 |
| Qfrt Y4Cys | Q frt | yes | yes | Y4 Cys | RC::FLTG | yes | 5 | 1/5 |
| frt Y4Linear | frt | yes | yes | Y4 Linear | RC::FLTG | no | 5 | 0/5 |
| frt Y4L5 | frt | yes | yes | Y4 L5 | RC::FLTG | no | 5 | 0/5 |
| noguidesY4Linear | no | yes | yes | Y4 Linear | RC::FLTG | no | 5 | 0/5 |
| noguides Y4L5 | no | yes | yes | Y4 L5 | RC::FLTG | no | 5 | 0/5 |
| onlyguides RC::FLTG | Qfrt | yes | yes | no | RC::FLTG | no | 3 | 0/3 |
| onlyguides | Q | yes | yes | no | FVB enriched | no | 4 | 0/4 |
| Q Y4Linear nude | Q | yes | yes | Y4 Linear | NMRInude | yes | 8 | 6/8 |
| Q Y4L5 nude | Q | yes | yes | Y4 L5 | NMRInude | yes | 9 | 2/9 |

Abbreviations: Q: crRNA for Rb, Rbl1, Pten, and Trp53; Qfrt: crRNA for Rb, Rbl1, Pten, Trp53, and frt

**Fig. 3.**
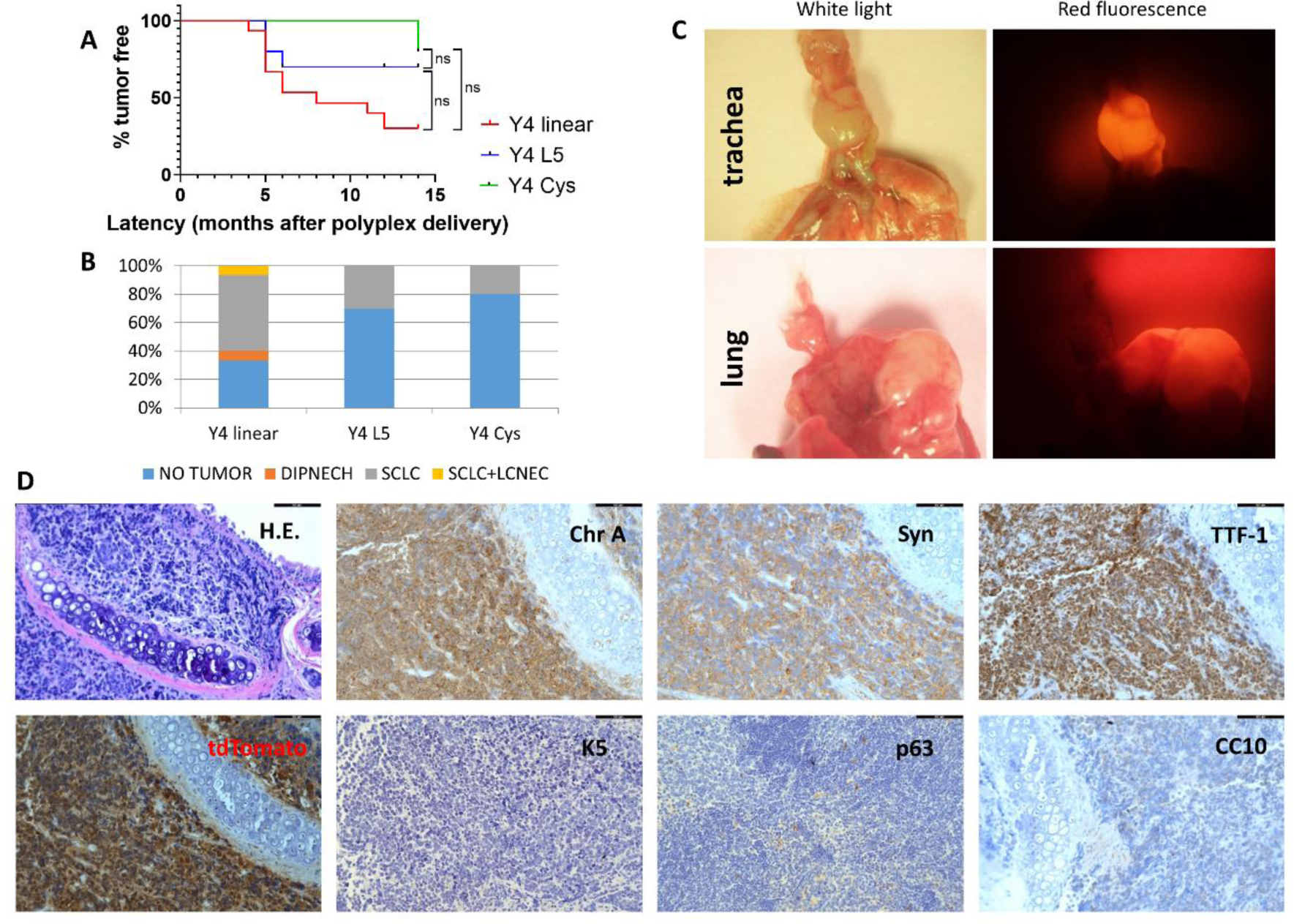
Gene editing of *Rb1, Rbl1, Pten*, and *Trp53* in adult RC::FLTG airway system leads to the development of Small Cell Lung Cancer (SCLC). A) Kaplan-Meier percent tumor-free survival curve of RC::FLTG mice administered with ribopolyplexes composed of Qfrt RNPs and different Y4 polymers family members: Y4 Linear (red line) Y4 L5 (blue line) or Y4 Cys (green line). X axis: latency (months after intratracheal delivery of the polyplexes). Total number of mice analyzed for Y4 linear, n = 15; Y4 L5, n = 10; Y4 Cys, n=5. *p<0.05 determined by Logrank test for trend B) Incidence and histopathology spectrum of tumors arisen from RC::FLTG mice administered with the polyplexes Qfrt Y4Linear (Y4 Linear), Qfrt Y4 L5 (Y4 L5), Qfrt Y4 Cys (Y4 Cys). Total number of tumors Qfrt Y4 Linear, n= 85; Qfrt Y4 L5, n=6; Qfrt Y4 Cys n=1. **C)** Stereomicroscope images of tumors in trachea (upper panel) or lung (lower panel) in RC::FLTG mice. Bright field (left) and fluorescence red light (right) images showing tdTomato detection. **D)** Representative images of a tracheal tumor. Hematoxylin–eosin (H.E.) and immunohistochemical analyses of the quoted proteins in a tracheal tumor. Bars = 100 μm. DIPNECH, Diffuse idiopathic pulmonary neuroendocrine cell hyperplasia; LCNEC, large-cell neuroendocrine carcinoma; SCLC, small-cell neuroendocrine carcinoma; ChrA, Chromogranin A; Syn, synaptphysin; TTF-1, Thyroid Transcription Factor-1; CC10 Clara/club cell secretory protein; K5, keratin 5; CC10 Clara/club cell secretory protein.

To test whether potential carcinogenic in vivo effect could result from any of the different components of the ribopolyplexes (RNP+ Y4polymers) separately, we set up controls including ribopolyplexes without gRNAs, RNPs without polymer, both in RC::FLTG and FVB enriched genetic backgrounds, and ribopolyplexes for *frt* guide only. The different experimental groups are detailed in Table 1. Tumors developed exclusively in mice inoculated with RNPs with guides for all four tumor suppressor genes plus polymers. None of the mice from control groups exhibited any signs of illness or abnormal signals during the CT scan follow-up. At 14 months post-inoculation, no tumors were observed in any control mice during necropsy (Table 1). Since no tumors developed when any of the ribopolyplex components were inoculated separately, gene editing events are the presumed cause of carcinogenesis.

Primary tumors were localized along the respiratory system (lungs and trachea), and were particularly abundant in trachea (Fig. 3C, Table S1). This could reflect the initial allocation of the ribopolyplexes along the respiratory system after intratracheal delivery. Histopathological analysis revealed that virtually all tumors were characterized by small-size cells with scant cytoplasm, inconspicuous nucleoli and nuclear molding (Fig. 3D. Fig. S7A). This histopathology is very similar to that of tumors previously obtained by conventional transgenesis and to human SCLC [6,8,35]. The diagnosis was further confirmed with immunohistochemical expression of neuroendocrine markers Synaptophysin, Chromoganin A, and TTF1 and absence of non-neuroendocrine markers such as CC10, Keratin K5 and p63 (Fig. 3D). Occasionally, Large Cell Neuroendocrine Carcinoma (LCNEC) and Diffuse Idiopathic Pulmonary Neuroendocrine Cell Hyperplasia (DIPNECH) were also observed in the *Qfrt Y4 Linear* group (Fig. 3B). These data show that the disruption of the tumor suppressor genes *Rb1, Rbl1, Pten* and *Trp53* in adult lungs by means of *in vivo* gene editing leads to the development of SCLC.

Tumors were dissected and DNA extracted for PCR/Sanger analysis to study the sequences of the regions where we wanted to introduce mutations in each of the four tumor suppressor genes: *Rb1, Rbl1, Pten* and *Trp53*. Out of 20 tumors analyzed, we found indels at the calculated positions in all tumors for the four TSGs (Fig. 4 and Supplementary Fig5), except for *Trp53* in one case (mouse 4nu) and *Rb1* in another (mouse 153) (Supplementary Fig5). Most mutations were frameshift, which was consistent with the generation of loss of function alleles. Multiple tumors arose in the respiratory tract of most mice (Table S1); Fig. 4 illustrates the indels found at the four tumor suppressor genes (TSGs) in three different tumors dissected from a single mouse. The mutations observed in each tumor are distinct combinations, highlighting that each tumor is likely the result of different mutagenesis events. However, as depicted in Fig. S5, not all tumors showed the presence of the tomato marker (highlighting the difference between passenger versus driver mutations). This underscores the intrinsic requirements of the participation of the four driver mutations in the SCLC oncogenic process.

**Fig. 4.**
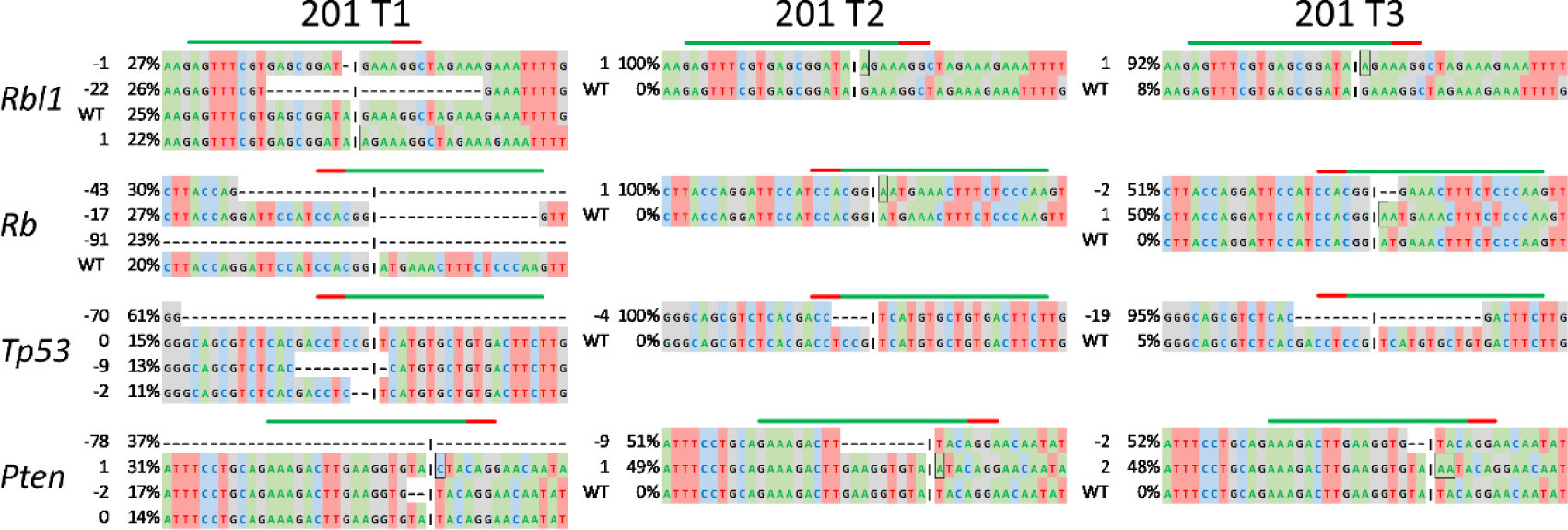
Tumor genotyping by Sanger Sequencing. Sanger sequences of PCR amplicons of the four tumor suppressor genes corresponding to 3 different tumors from one animal (mouse 201). T: tumor. Green line: gRNA target sequence; Red line: PAM sequence. Indel length and percentage shown on the left. Black rectangle boxes depict insertions. Cleavage site indicated by black vertical lines.

In summary, these data show that directing genome editing to lung cells in adult mice using polyplexes is a viable method to trigger the spontaneous formation of tumors when tumor suppressor genes are targeted. Furthermore, it’s important to note that the polyplex reagents themselves do not possess inherent tumorigenic properties. (Table 1).

### Transcriptomic analysis of gene editing mediated SCLC tumors

To further characterize our models, we performed RNA sequencing from seven mouse SCLC samples (tumor group) and five normal lung tissue samples from littermates (control group). As expected, principal component analysis (PCA) and hierarchical clustering (HCL) showed that samples from each group cluster together (Fig. 5A, B). Importantly, tumor and control samples showed clear differences in the bidimensional PCA space (Fig. 5A) or patterns of differentially expressed genes (Fig. 5B), which demonstrated strong differences in transcriptome programs. To further demonstrate that *in vivo* gene editing-generated tumors are SCLC, we performed gene set enrichment analysis (GSEA) using reported signatures from mouse and human SCLC. Tumors resulting from *in vivo* gene editing are significantly enriched in genes that we previously reported to be overexpressed in K5-QKO SCLC tumors (moSCLC signature of 1335 genes)[6] (Fig. 5C). Our gene editing-generated mouse tumors are also significantly enriched in genes used to classify human SCLC (Class IV signature) [1] (Fig. 5D). Altogether, our RNA-seq analysis demonstrates that tumors obtained with our *in vivo* gene editing strategy display the gene expression program of mouse and human SCLC.

**Fig. 5.**
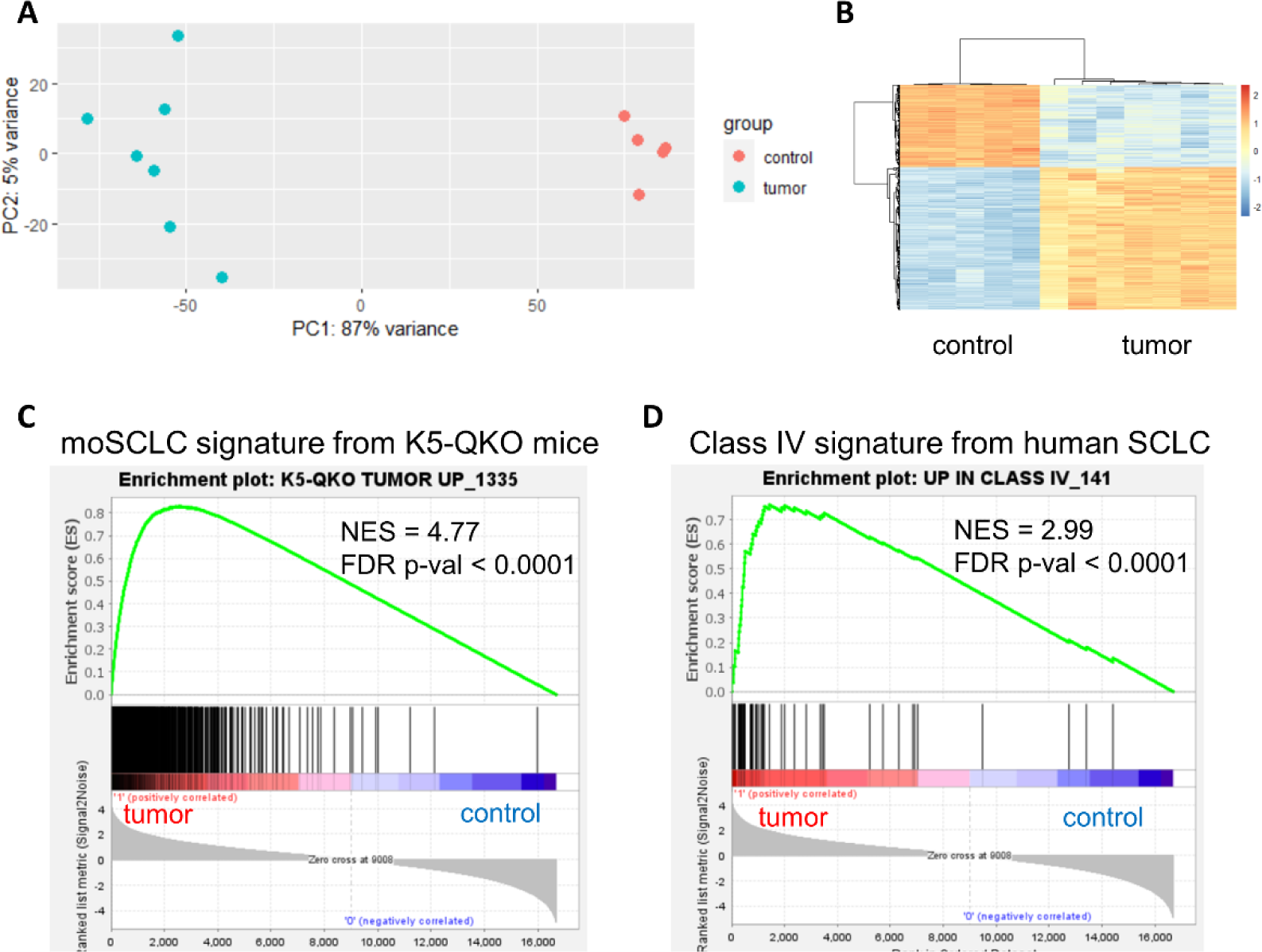
Transcriptome analysis of SCLC tumors developed by gene editing. Principal component analysis (PCA) (A) and hierarchical clustering (HCL) (B) showed that replicates from each sample type (tumor or control samples) are grouped together, highlighting clear differences between tumor and control samples. Controls include five normal lung samples, and tumors include seven lung cancer samples with histopathological features of SCLC. C) Gene set enrichment analysis (GSEA) plots showing enrichment in the tumors in moSCLC gene signatures established in the K5-QKO mouse model. Note that the positions of the moSCLC genes are skewed toward the left end of the rank-sorted list, reflecting their statistically significant induction in tumors. D) GSEA analysis showing enrichment in gene editing-generated mouse tumors in a signature from human SCLC.

### Lung carcinogenesis by gene editing in immunodeficient mice

One of the challenges of germline genetic manipulation is the extraordinary difficulty involved in creating genetically modified mouse models in pure genetic backgrounds. The process outlined here has the added advantage of being adaptable to any genetic background, including immunodeficient strains. We administered polyplexes to immunodeficient NMRI nu/nu (nude) mice to recapitulate the carcinogenesis procedure described above for an immunocompetent background (Table 1). This experiment also allowed the study of the possible effect of immunodeficiency on the process of lung carcinogenesis.

We treated 17 mice with quadruple (cr RNA *for Rb1, Rbl1, Pten* and *Trp53*) RNP complexes (treatment group designated as Q). Eight mice were treated with Y4 Linear polymers (“*Q Y4Linear nude*”) and 9 mice with Y4 L5 polymers (“*Q Y4L5 nude*”, Table 1; Fig. 5). Nude mice developed lung tumors with a latency of 4–12 months (*Q Y4Linear nude*) or 7–8 months (*Q Y4 L5 nude*) with an incidence of 75% and 24%, respectively (Fig. 5A, B). Kaplan-Meier survival curve was significantly different between the Y4 Linear and Y4 L5 treated groups (log-rank p < 0.01) (Fig. 5A). No significant differences in survival (determined by log-rank test) were observed between immunodeficient or immunocompetent mice treated with the same polymer either for Y4 linear or Y4 L5 polyplex treatments (Fig. S5). Following the same pattern as in the immunocompetent background, tumors were diagnosed as SCLC, and occasionally as LCNEC (Fig. 6B, Table S1). These findings were further confirmed by immunohistochemical analysis of markers (positive expression of CGRP, Syn, TTF1 and absence of expression of CC10) (Fig. 6F). Tumors were frequently seen in trachea, likely evolving from keratin K5 positive epithelial basal cells, although no expression of this keratin was retained in the tumor, as previously observed (Fig. 6 G) [6].

**Fig. 6:**
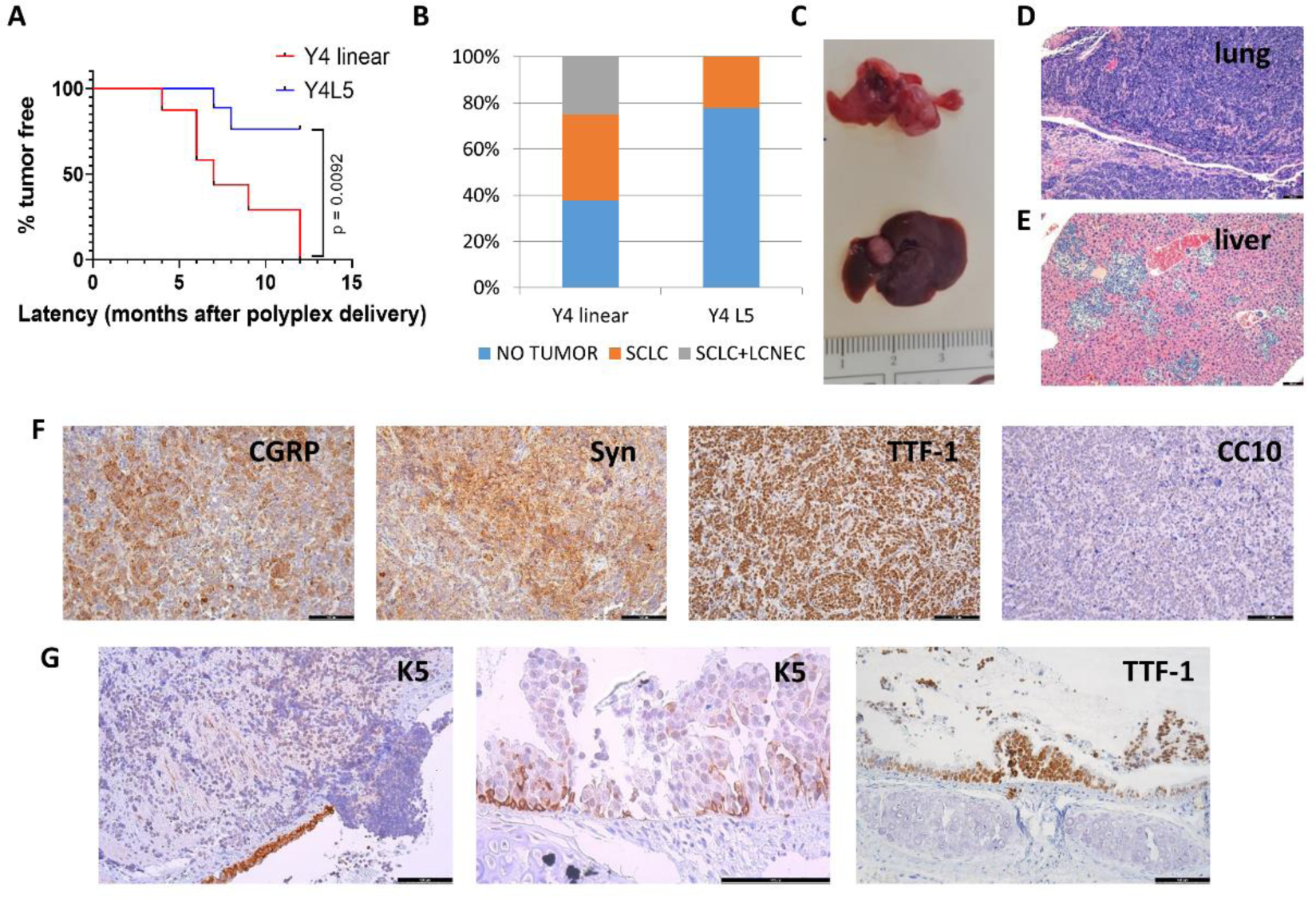
Lung carcinogenesis by gene editing in immunodeficient mice. **A)** Kaplan-Meier tumor-free survival curve of *Q Y4Linear nude* (red line n=8) and *Q Y4L5 nude* (blue line n=9) mice ** *p* ≤ 0.01 determined by log-Rank. x axis: latency (months after inoculation starting point). **B)** Incidence and histopathology spectrum of tumors arisen in nude mice administered with the polyplexes QY4 Linear (Y4 Linear), QY4 L5 (Y4 L5). Total number of tumors Q Y4 Linear nude (n= 59); Q Y4 L5 nude (n= 11) **C)** Gross appearance of tumors in lung (up) and liver (down). **D)** Hematoxylin–eosin (H.E.) of a representative lung SCLC and **E)** liver metastases. **F)** Immunohistochemistry staining of lung tumors as quoted. **G)** Tracheal tumor evolving from K5 expressing basal cells. Bars= 100 µm. CGRP, Calcitonin Gene Related Protein, Syn, synaptophysin; TTF-1, Thyroid Transcription Factor-1; CC10 Clara/club cell secretory protein; K5, keratin 5.

These data demonstrate that the procedure described in this study enables the creation of complex lung cancer mouse models in various genetic backgrounds, including in strains that are complex and troublesome to manipulate such as immunodeficient mice.

### Development of metastasis in non-viral gene edited SCLC tumor bearing mice

SCLC is a highly metastatic carcinoma, with reported metastatic rates about 70% in human patients at diagnosis [36]. Thus, we performed complete analyses in search of metastases. Five out of 10 tumor-bearing mice in the *Qfrt Y4 Linear* group had metastases in liver and/or lymph nodes (Table S1) that displayed histological features and neuroendocrine marker expression similar to those in primary lung tumors (Fig. S5). These features were also described in the tumors previously generated with the cre/loxP system [6] and resemble the metastatic behavior of human SCLC.

To characterize metastases, we performed expression analysis of diagnostic markers of SCLC. As we found for primary tumors the metastases expressed CGRP, Synaptophysin and TTF-1 and were negative for CC10. Tomato expression could be observed only in some primary tumors (Fig. S8) and their corresponding metastases (Fig. S8). High-grade neuroendocrine lung tumors SCLC (and LCNEC) arised as a consequence of simultaneous gene editing of the 4 tumor suppressor genes (Fig. 4, Fig. S5) but are not necessarily linked to frt gene editing.

Liver and lymph node metastases were also found in immunodeficient mice, with higher frequency and metastasis burden than in the immunocompetent background (4 out of 6 tumor-bearing mice in the *Q Y4 Linear nude* group and 1 out of 2 in the *Q Y4L5 nude* group)(Table S1). The larger size of the liver metastases in the nude background allowed us to readily isolate their DNA and perform sequencing analysis in combination with primary tumors (Fig. S7). Tumor genotyping showed that the tumor suppressor genes carried indels at the expected sites. We observed that three metastatic tumors had identical mutations to one of the primary tumors in one animal (mouse 10nu) and one metastasis had identical mutations to a primary tumor in another animal (mouse 4nu) (Fig. 6 and Fig S5).

Taken together, these results show that respiratory tract cell-targeted genome editing in adult mice using a cationic polymer-based non-viral delivery approach is sufficient to induce spontaneous tumor formation. The set of targeted genes determines the tumor to be developed (disruption of *Rb1 Rbl1, pTen and Trp53* genes leads to generation of SCLC irrespective of the methodology used). In conclusion, our CRISPR/Cas9 RNP delivery system utilizing cationic polymers offers an efficient approach for modeling lung tumorigenesis by simultaneously inactivating a set of tumor suppressor genes,which will enable rapid and cost-effective hypothesis testing independent of the mouse genetic background and without the need for germline manipulation or custom viral vectors.

## DISCUSSION

Lung cancer mouse models have provided insight for cancer research and advances in patient care [37–40]. However, these models were mainly based on complex genetic engineering modifications that are extremely time-consuming, labor-intensive and involve numerous animals. In this study, we have developed lung carcinogenesis models that circumvent the necessity for individually engineered alleles for each candidate gene to be characterized or to construct viral vectors for the delivery of gene editing tools, which considerably reduces the number of animals required. Our efforts have been directed at establishing an *in vivo* CRISPR delivery system based on nanoparticles formed by CRISPR RNPs with non-viral chemical carriers. These carriers are linear and newly synthesized branched PBAE polymers. In combination with CRISPR/Cas9 RNPs they form nanoparticles called ribopolyplexes. In a previous study we demonstrated the ability of PBAE ribopolyplexes to accomplish gene editing using cultured human keratinocytes, suggesting the potential therapeutic utility of these reagents for the correction of genetic pathologies [29]. Here, we present evidence supporting the feasibility of introducing gene modifications in somatic cells of the adult respiratory tract through gene editing using CRISPR/Cas9 ribopolyplexes. This approach simultaneously targets multiple tumor suppressor genes, which resultis in tumors carrying mutations in these specific genes.

*In vivo* gene editing in the lung with CRISPR has been previously demonstrated using adenoviral [41] and AAV [42] vectors capable of activating reporter alleles upon editing with guides that remove transcriptional STOP sequences. Here, we use a new non-viral carrier system based on CRISPR ribopolyplexes to achieve levels of editing comparable to those obtained with viral vectors. To assess and demonstrate *in vivo* gene editing, we have established a new model for the activation of a reporter allele of tdTomato protein expression following gene editing by CRISPR using a guide targeting the frt sequence. In this model system we find that ribopolyplexes produce gene editing in a variety of cell types, which points to the utility of these carriers to modify respiratory tract tumor-initiating cells. Other non-viral carriers based on amphiphilic peptides have been developed to deliver CRISPR/Cas9 into airway tissue, thus demonstrating their efficiency in activating reporter alleles [43,44]. In addition, gene editing in the lung, which was demonstrated through the use of a reporter allele, has also been achieved using LNPs as carriers of CRISPR RNPs after intravenous injection [20]. Recent advances in the delivery of CRISPR to the lung as RNAs in LNP formulations [45] suggest that this approach will also be useful for somatic mutagenesis of cancer related genes in this organ. Taken together, these findings show progress toward novel chemical carrier-based strategies for lung cancer modeling. In addition, the development of recombinant gene editing enzymes based on Cas9 fusion proteins with extended capabilities like base editing [46] will likely open up new avenues for precise modifications with CRISPR/Cas9 RNPs, which will include the ability to engineer activating mutations in oncogenes alongside loss-of-function mutations. Indeed, the activation of oncogenes in lung tumor modeling through HDR using chemical carriers has been described [20]. These findings suggest the emergence of potentially valuable additional tools for fast and flexible lung cancer modeling.

By using Cre/ loxP viral recombination strategies, we previously described how the ablation of four tumor suppressor genes (*Rb1, Rbl1, Pten, Trp53*) in the respiratory tract led to the development of high-grade neuroendocrine tumors (LCNEC and SCLC) that grew differentially depending on the targeted epithelial cell type [6,32]. In this study, by delivering CRISPR/Cas9 reagents as ribopolyplexes with PBAE polymers we demonstrate the feasibility of generating tumors by mutagenesis in mice without any prior gene manipulation. Histopathological traits, molecular characteristics and transcriptomic profiles showed that these SCLC tumors were similar to those generated with our Cre/loxP strategy [6] and human SCLCs. This innovative approach opens the possibility of using this platform to streamline the evaluation of candidate genes’ functionality in lung tumor development. In our study, we simultaneously disrupted four tumor suppressor genes that are known to be associated with SCLC development. Adding a fifth gRNA to target the frt sequence in the RC::FLTG mice experiment led to highly visible red fluorescent tumors, making it easier to spot neoplasic lesions caused by gene editing. This demonstrates that it is possible to simultaneously modify at least five genes, and it underscores that gene editing can modify genes unrelated to tumor formation when a combination of genes related to tumorigenesis and non-tumor-causing genes are targeted.

In our previous work, we observed that ablation of these four suppressor genes by adenoviral vectors with Cre recombinase expression, targeted by the ubiquitous CMV promoter to a wide variety of epithelial cell types, predominantly led to the formation of LCNECs. In contrast, when Cre expression was targeted to basal cells of the respiratory epithelium located in the trachea and bronchi by the keratin 5 promoter, tumors were predominantly SCLC [6,32]. SCLC can originate from several cell types in the lung, including CGRP+ neuroendocrine cells, SPC+ type II alveolar cells, CC10+ alveolar type I and K14+ basal cells [7,47,48]. The tumors generated by ribopolyplex-mediated gene editing were mostly SCLC. This observation raises questions regarding the preferential targeting of K5+ cells in our experimental conditions, potentially explaining the significant occurrence of tumors within the trachea, where basal K5+ cells are predominantly found. Differences in the physical and chemical characteristics of the reagents could account for distribution pattern variations in different respiratory tract locations when compared to viral vector-based methods. In any case, the homogeneity observed in the appearance of SCLC tumors in this study may offer advantages for preclinical drug testing.

Considering the varying susceptibility to carcinogenesis among different mouse strains, the genetic background in which carcinogenesis models are developed is an important consideration [49–51]. Having a technology that facilitates the introduction of a set of mutations with ease, regardless of the genetic background, is highly advantageous. When we transferred our gene suppressor tumor disruption experiment to immunodeficient nude mice, which have a genetic background less amenable to gene manipulation, we confirmed that the carcinogenesis works in a manner comparable to that found in the RC::FLTG mice (C57Bl/6J background [31]). Treated nude mice developed liver metastases, with both primary tumors and metastases harboring mutations at the intended locations of tumor suppressor genes. The mutations in the metastases mirrored those found in one of the primary tumors. Immunocompetent mice also developed liver metastases, but these were larger and more numerous in the immunodeficient model. Given the straightforward extrapolation of the protocol, its implementation in animal models other than mice, such as rats, is conceivable through the adaptation of gRNA sequences.

We tested three related polymers in this study: Y4 Linear, previously described [25], and two branched variants, Y4 L5 and Y4 Cys, which differ in their end-capping. Tumors developed with all three variants, with the linear polymer demonstrating slightly greater efficiency, and only one tumor developing in the cysteine-modified variant. Although the efficiency of in vivo gene editing in lung tissue is limited, as deduced from the expression of the tdTomato protein in isolated clusters of cells distributed throughout the lung tissue, the occurrence of tumors in a significant percentage of the treated mice suggests that the incidence of mutagenesis is appropriate for modeling the natural somatic mutation frequency underlying tumor development.

This study presents a potentially more straightforward and cost-effective alternative for developing lung cancer models across various mouse genetic backgrounds. The ease of implementing mutagenesis using a chemical carrier platform with ready-to-use synthetic reagents opens the door to designing functional screens for testing candidate cancer-related genes in combination with known carcinogenic genes. As gene editing technology continues to advance, the potential modifications achievable through chemical carrier delivery of CRISPR RNPs will likely extend beyond gene inactivation via the introduction of indels, thereby expanding the scope for cancer modeling based on non-viral carriers platforms.

## MATERIALS AND METHODS

### -Mice/Animal Studies

All the animal work was approved by the Animal Ethical Committee (*CEEA*) and conducted in compliance with *Centro de Investigaciones Energéticas, Medioambientales y Tecnológicas* (CIEMAT) guidelines. Specific procedures were approved by *Comunidad Autónoma de* Madrid (protocol code ProEX 208/15, date of approval 13 July 2015, ProEX 111.1/21, date of approval 5 March 2021 and ProEX 0730/22, date of approval 25 May 2022).

RC::FLTG (Plummer 2015) and C57BL/6J mice were obtained from Jackson (Jax 026932 and Jax 2498079-86 respectively, Charles River) and then bred in our facilities. The FVB enriched mice used have been described elsewhere [9]. NMRI-Foxn1 ^nu/nu^ immunodeficient 6-8 week old mice were obtained from Janvier (Saint-Berthevin).

Animals were inoculated with a single application of 50-100 µL of CRISPR/Cas RNP ribopolyplexes (or control reagents, see Table 1) at 6-12 weeks of age by intratracheal intubation as described [5].

### -Cell culture and nucleofection

#### MEF isolation

Freshly harvested 13.5 d.p.c. embryos were placed in a cell culture dish and minced with a razor blade after the addition of trypsin. The dish was incubated at 37°C for 30-45 minutes, and trypsin activity was neutralized by adding DMEM 10% FCS medium. The tissue was further disrupted by pipetting, and the cell suspension was transferred to a larger flask or plate containing medium. The cells were then grown to confluence for 3-4 days, trypsinized and the suspension transferred to larger flasks. The cells were again grown to confluence for 3-4 days. Finally, the cells were harvested using trypsin and frozen in vials at a concentration of 3×106 cells per vial.

Nucleofection of CRISPR/Cas9 RNPs in MEFs was performed as previously described (Bonafont et al, PMID: 30930113).

#### Editing efficiency analysis by Cytometry

MEFs nucleofected with CRISPR/Cas9 RNPs with gFrt guide were analyzed by flow cytometry in a LSR Fortessa (BD) cytometer for red fluorescence.

### -Polymer synthesis, characterization and nanoparticle formation

#### Polymer synthesis

Y4 Linear was synthesized by copolymerizing 5-amino-1-pentanol (S5, A2 type monomer) and 1,4-butanediol diacrylate (BDA, C2 type monomer) [C32, [25]. For the branched version, ethane-1,2-diamine (EDA, B4 type polymer) was added to BDA and S5 via a facile one-pot “A2 + B4 + C2” Michael addition reaction. The ratio of amines to acrylate was set at 1.2: 1. The molecular weight (Mw) of the polymers was monitored with gel permeation chromatography (GPC) and reactions were stopped when the polymers reached the desired molecular weight by cooling the reactions to room temperature. Then, 1,3-diaminopropane (103) diamine was used to end-cap the excessive acrylate groups in Y4 Linear and Y4 L5 by adding it at room temperature for 24 h. Cysteine was used instead of 103 for Y4 Cys.

To purify the polymer, it was precipitated three times using an excess of diethyl ether and subsequently dried in a vacuum oven for 24 hours [27]. Monomers were all purchased from Merck (Ireland). Polymers were dissolved in DMSO to 100 μg/μL stock solution and stored at - 20 °C.

#### Polymer characterization (GPC and NMR)

Polymer samples were diluted in dimethylformamide 5 mg/mL (DMF), and after filtration (0.22 μm filter) were injected into a gel permeation chromatographer, (1260 Infinity MDS GPC System, Agilent Technologies, Ireland) equipped with a refractive index detector (PI), a viscometer detector (VS DP) and a dual angle light scattering detector (LS 15° and 90°) for molecular weight sample determination. Columns were eluted with DMF with 0.1% LiBr as a solvent at a flow rate of 1 mL/min and at 60 °C. Poly(methyl methacrylate) (PMMA) standards were used for calibration. To monitor the molecular weight of polymers during polymerization, a volume of 50 μL of the reaction mixture was analyzed. To confirm chemical structure, composition and purity by proton nuclear magnetic resonance (^1^H NMR), polymer samples were dissolved in CDCl_3_ and NMR measurements were conducted on a Varian INOVA 400 MHz Spectrometer (Edinburgh, United Kingdom).

#### Ribopolyplex formation

Nuclease-free duplex buffer was used to dilute crRNAs and tracrRNA to 200 µM and mixed following manufacturer recommendations with HiFi Cas9 nuclease (IDT, Coralville, IA, USA) at sgRNA(crRNA+tracrRNA):Cas9 molar ratio 6.6:1.

Nanoparticles were formed by complexation of the CRISPR/Cas9 RNPs with the polymers at 20:1 weight/weight ratio (polymer:RNP). Each polymer (100μg/μL in DMSO) and CRISPR/Cas9 RNP complex, with the corresponding gRNA, was dissolved in 25 mM sodium acetate (SA, Sigma Aldrich, St. Louis, MO, USA) at 1:1 volume/volume in a total volume of 100 μL. The dissolved polymer was added dropwise to the dissolved RNP complex, mixed by vortexing for 30 seconds and incubated for 15 mins at room temperature.

### -CRISPR/Cas9 gRNA design

gRNAs were designed using a CRISPR design online tool (http://crispor.tefor.net/crispor.py). Synthetic RNAs and recombinant Cas9 were purchased from Integrated DNA Technologies (IDT, IL), and RNP complexes were reconstituted according to the manufacturer’s instructions **gRNA sequences were:** gfrt-1: 5**’-**TCCTATTCTCTAGAAAGTAT-3’ (fw); gfrt-2: 5’-CTATACTTTCTAGAGAAT-3’ (rev); p107-E3: 5’-AGTTTCGTGAGCGGATAGAA-3’ (Exon 3, fw); Pten : 5’-AAAGACTTGAAGGTGTATAC-3’; Rb1-E2-5’: 5’-TTGGGAGAAAGTTTCATCCG-3’; p53 Walton-E5: 5’-GAAGTCACAGCACATGACGG-3.

### -Genotyping of gene-edited cells and tumors

Genomic DNA was isolated by isopropanol precipitation of cells and tumor lysates (lysis buffer was Tris [pH 8] 100 mM, EDTA 5 mM, SDS 0.2%, NaCl 200 mM, and 1 mg/mL proteinase K [Roche Diagnostics, Mannheim, Germany]) and resuspended in Tris/EDTA (TE) buffer. Approximately 20–50 ng genomic DNA was used for PCR amplification. PCR fragments spanning the nuclease target sites were generated with primers: Pten F : 5’ -GTGAGTGGCTGACTGTCCAG-3’; Pten R: 5’-GTGAGAGCTGTCAGGTGCTT-3’; Trp53-F: 5’-CCACCTTGACACCTGATCG-3’;Trp53-R: 5’- TAGCACTCAGGAGGGTGAG-3’; Rb1 F 5’-GCCAGTTCAATGGTTGTGGG-3’;Rb1 R3’-TGTCAGAGAAAGAGCTTGGCT-5’;Rbl1 F 5’-GCAAGATGCACGCAACAATC-3’;Rbl R 3’-TGGACATGTCAAACCTACCACA-5

### -Sequencing

Regions targeted for gene editing were PCR-amplified and Sanger sequenced. PCR products were treated with Illustra ExoProStar (GE Healthcare, UK), sequenced using Big Dye Terminator version (v.)1.1 Cycle Sequencing kit (Thermo Fisher Scientific, Waltham, MA) and examined on a 3730 DNA Analyzer (Life Technologies, Carlsbad, CA). Sanger traces were analyzed for indel deconvolution with the TIDE (http://tide.nki.nl) and DECODR (https://decodr.org/) webtools.

### -Computed Tomography (CT) imaging

Tumor development monitoring was performed by Computed Tomography (CT) imaging every 2 months after intratracheal inoculation, in a small-animal Super Argus 3r PET/CT scanner (Sedecal, Spain) with the following acquisition parameters: 500 μA, 30 kV, 360 projections, 1 shot and 4×4 Binning

### -Histology and immunostaining

At necropsy, tumors from RC::FLTG mice were examined for red fluorescence emitted by the protein reporter tdTomato under a magnifying glass equipped with white and green light. Lungs were perfused with 4% formaldehyde. Samples were fixed in 4% buffered formalin and embedded in paraffin wax. Sections (5 μm) were stained with hematoxylin and eosin (H/E) for histological analysis or processed for immunostaining.

#### Immunohistochemistry

Immunohistochemical analyses were performed essentially as in previously described standard protocols [33,52,53]. Leica microscope was used and images obtained with LAS X software (v3.7.4, Leica Microsystems, Wetzlar, Germany). Primary antibodies were used as follows: anti-tdTomato (NB-25-00005, NeoBiotech, Nanterre, France, 1:300); anti-mCherry (ab167453, Abcam, Cambridge, UK, 1:300); anti-chromogranin A (ab15160, Abcam, 1:100); anti-TTF1 (ab76013, Abcam, 1:200); anti-p63 (ab53039-100, Abcam, 1:50); anti-CGRP (c8198, Merck, Burlington, MA, USA, 1:2000); anti-Keratin 5 (PRB-160P, BioLegend, Dedham, MA, USA, 1:500); anti-CC10 H-75 (sc-25554, Santa Cruz Biotechnology, Dallas, TX, USA, 1:300); anti-synaptophysin SY38 (ab8049, Abcam, 1:100); anti-RB (554136, BD Biosciences, Franklin Lakes, NJ, USA, 1:100); anti-p107 (sc-250, Santa Cruz Biotechnology, 1:100); anti-p53 (NCL-L-p53-CM5p, Leica Biosystems, Wetzlar, Germany, 1:200); anti-PTEN (9559, Cell Signaling Technology, Danvers, MA, USA, 1:200). Secondary antibodies for immunohistochemistry were used as follows: biotin anti-rabbit (No. 711-065-152, Jackson ImmunoResearch, West Grove, PA, USA, 1:1000); biotin anti-goat (No. 705-065-147, Jackson ImmunoResearch, 1:1000); biotin anti-mouse (No. 715-065-151, Jackson ImmunoResearch, 1:1000).

#### Immunofluorescence

Tissues were fixed in 4% buffered formalin for 24 h and afterwards dehydrated and embedded in paraffin wax. Sections (5 μm) were used for immunofluorescence analyses, performed as follows: the sections were (1) deparaffinized; (2) incubated with 10% horse serum for 30 min at 37 °C to block non-specific binding; and (3) washed three times with sterile phosphate-buffered saline (PBS) (pH 7.5) prior to incubation with the appropriate primary antibodies diluted in horse serum 10%. Primary antibodies were used as follows: 1/200 anti-RFP (MBS448122, MyBioSource, Vancouver, Canada); 1/300 dilution of anti-tdTomato (NB-25-00005, NeoBiotech); 1/300 dilution of anti-mCherry (ab167453, Abcam); 1/2000 dilution of anti-CGRP (c8198, Merck, Burlington, MA, USA); 1/500 dilution of anti-Keratin 5 (PRB-160P, BioLegend, Dedham, MA, USA); 1/300 dilution of anti-CC10 T-18 (sc-9772, Santa Cruz Biotechnology, Dallas, TX, USA); 1/300 dilution of anti-CC10 H-75 (sc-25554, Santa Cruz Biotechnology); 1/300 dilution of anti-Aquaporin 5 (ab78486, Abcam) and 1/1000 dilution of anti-proSPC (AB3786, MilliporeSigma, Merck, Burlington, MA, USA).

An additional step prior to secondary antibody incubation was followed if quenching autofluorescence was necessary (lung parenchyma staining): (4) 30 min incubation in ammonium chloride (NH_4_Cl) and (5) 4 min incubation in Tissue Autofluorescence Quenching Kit (ReadyProbes, ThermoFisher Scientific).Secondary antibodies were used as follows: 1/1000 dilution of anti-rabbit AlexaFluor488 (A-21206, Invitrogen, Thermo Fisher Scientific, Waltham, MA, USA); 1/1000 dilution of anti-rabbit DyLight594 (SA5-10040, Invitrogen); 1/1000 dilution of anti-goat AlexaFluor488 (A-11055, Invitrogen) and 1/1000 dilution of anti-goat AlexaFluor647 (A-21447, Invitrogen). Diamidinophenylindole (DAPI) was used to counterstain the nuclei or chromosomes.

#### Dual color immunohistochemistry

Double immunohistochemistry was performed as sequential double chromogenic IHC using BOND RX fully automated research stainer (Leica Biosystems, Wetzlar, Germany) with different detection systems based on polymers linked to goat polyclonal antibodies against rabbit IgGs: The chromogen bond polymer detection systems where: BOND Polymer Refine DAB detection, BOND Polymer Refine Red Detection (which is a biotin-free IVD labeled red detection system, that utilizes alkaline phosphatase (AP)-linked compact polymers) and a BOND Polymer HRP PLEX detection with a Green chromogen. Primary antibodies were used as follows: anti-mCherry (ab167453, Abcam, 1:1000); anti-CGRP (c8198, Merck, 1:2000); anti-Keratin 5 (PRB-160P, BioLegend, 1:2000); anti-CC10 H-75 (sc-25554, Santa Cruz Biotechnology, 1:1000); anti-CGRP (c8198, Merck, 1:2000); anti-Aquaporin 5 (ab78486, Abcam, 1:2000); anti-proSPC (AB3786, MilliporeSigma, 1:4000).

### RNA sequencing

Tissues were embedded in RNALater (Ambion Inc., Thermo Fisher Scientific, Waltham, MA, USA) and RNA was isolated and purified using miRNeasy Mini Kit (Qiagen) according to the manufacturer’s instructions. RNA yield and quality were determined using an Agilent 2100 Bioanalyzer. Transcriptome sequencing was performed from n = 5 lung control littermates and n=7 mouse tumors. RNA sequencing was performed after poly-A selection using Illumina NovaSeq 2 x 150 bp sequencing. Reads were aligned to GRCm39 mouse genome with “Rbowtie2”, counts extracted with featureCounts function from “Rsubread” and gencode annotation release 27 (gencode.vM27.annotation.gtf), and differential expression with DESeq2. Principal component analysis graph was plotted after variance stabilizing transformation (vst function in DESeq2).

### Gene set enrichment analysis (GSEA)

We developed a signature of genes specifically up-regulated in tumors from our K5-QKO SCLC (moSCLC) models [6,33]. Briefly, the Affymetrix probe set identifiers for genes significantly up-regulated (FDR < 0.01; >2-fold) in moSCLC versus normal lung (1335 Affy IDs), and these were manually curated to generate a gene signature: moLCNEC signature (Supplementary Dataset S1). Gene set enrichment analysis (GSEA, www.broadinstitute.org/gsea (accessed on 22 sept 2022)) [54] was used to analyze enrichment of these gene signatures in the mouse tumors.

Similarly, enrichment was performed using the mouse orthologs of the Class IV signature of genes able to classify human SCLC (Supplementary Dataset S1) [1]. GSEA was performed from normalized counts after removal of genes with counts = 0 in all samples.

All the transcriptome-sequencing data are available at Gene Expression Omnibus ((GEO) http://www.ncbi.nlm.nih.gov/geo/, accession number GSE249293

### Data and Statistical Analyses

Data were analyzed with GraphPad Prism 9 software (GraphPad Software, La Jolla, CA, USA). Statistical analysis was performed using Log-rank (Mantel-Cox) test.

## Acknowledgments

We are indebted to Iván Seller for his great contribution during the time he was working in the lab, the personnel of the Animal Facility at CIEMAT for excellent care of the animals, Pilar Hernández for excellent technical assistance, and Norman Feltz for copyreading the manuscript. Many thanks to Prof Wanǵs group members at the University College of Dublin, especially Xianqing Wang for the preliminary polymer screening *in vitro*.

This research was funded by *Asociación Española contra el Cancer* (AECC) through the project Seed Idea IDEAS211079SANT to M.S.; Instituto de Salud Carlos III (ISCIII) through the projects CB16/12/00228/CIBERONC, PI21/00764 to M. S., PI21/00171 to R. M., and PI15/00993 to M. S. and co-funded by FEDER and the European Union; Science Foundation Ireland (SFI) through the projects Future Innovator Prize (18/FIP/3576) to W. W., Frontiers for the Future 2019 call (19/FFP/6522) to W. W.; China Scholarship Council (CSC202008300033) to W. W., and project Garcia-Diez 1 supported by EB-Research Network funded by DEBRA Spain, DEBRA Austria and DEBRA Sweden. E.R was supported by a FEDER co-funded grant (ref. PEJ-2020-AI BMD-18428) from the *Comunidad de Madrid*.

## Author contributions

R.M. and M.S. design research; I.L.S., A.M., E.R., M.G., M.O., M.I.G., D.P-V., R.M. and M.S. performed research; I.L.S.,Y.L.,M. O.,S.A,W,W.,R.G-E.,R.M. and M.S, contributed new reagents/analytical tools; I.L.S.,A.M.,E.R.,Y.L.,M.G.,M. O.,M.I.G.,A.B..E, D.P-V., S.A, W.W., R.G-E., R.M. and M.S. analyzed the data; R.M. and M.S. wrote the paper

## SUPPLEMENTARY FIGURES

**Fig. S1.**
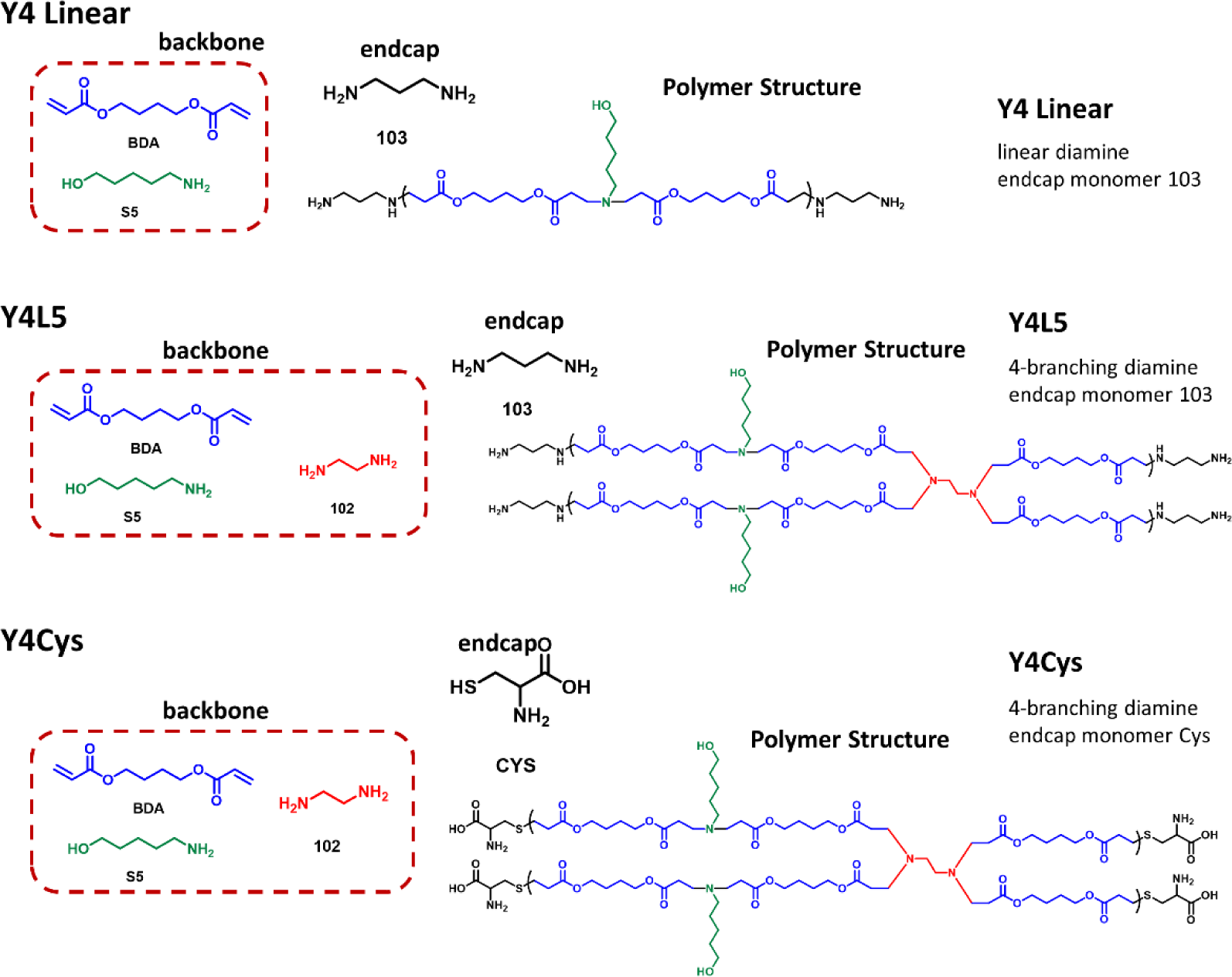
Chemical structure of the Y4 family of cationic polymers used as non-viral delivery vectors (Y4 Linear, Y4 L5, Y4 Cys). Backbone and end cap monomers that compose the Y4 Linear, Y4 L5 and Y4 Cys polymers. Y4 Linear and Y4 L5 (a 4 branched polymer generated by adding a 4-branching diamine to Y4 Linear) contain an endcap monomer 103 which confers an overall positive charge to the ribopolypexes to induce cellular uptake. Y4 Cys uses cysteine as an endcap monomer to confer both positive and negative charges to promote protein binding.

**Fig. S2.**
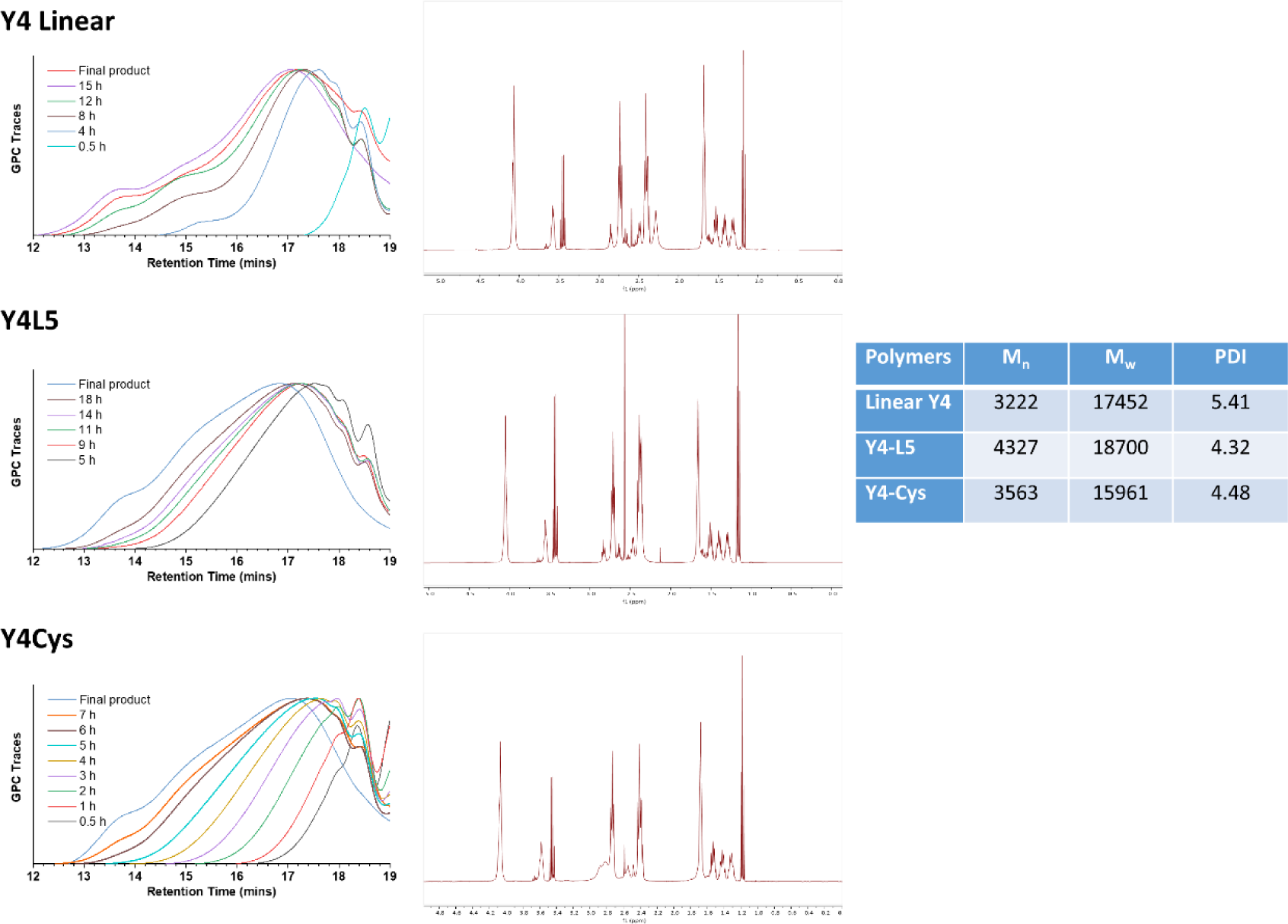
Characterization of the Y4 family of cationic polymers. Gel Permeation Chromatography (GPC, left) and Nuclear Magnetic Resonance (NMR, middle) traces for Y4 linear, Y4 L5 and Y4 Cys synthesis. Number average molecular weight (*M*_n_), average molecular weight (*M*_w_), and polydispersity Index (PDI) were determined by GPC analyses (right).

**Fig. S3.**
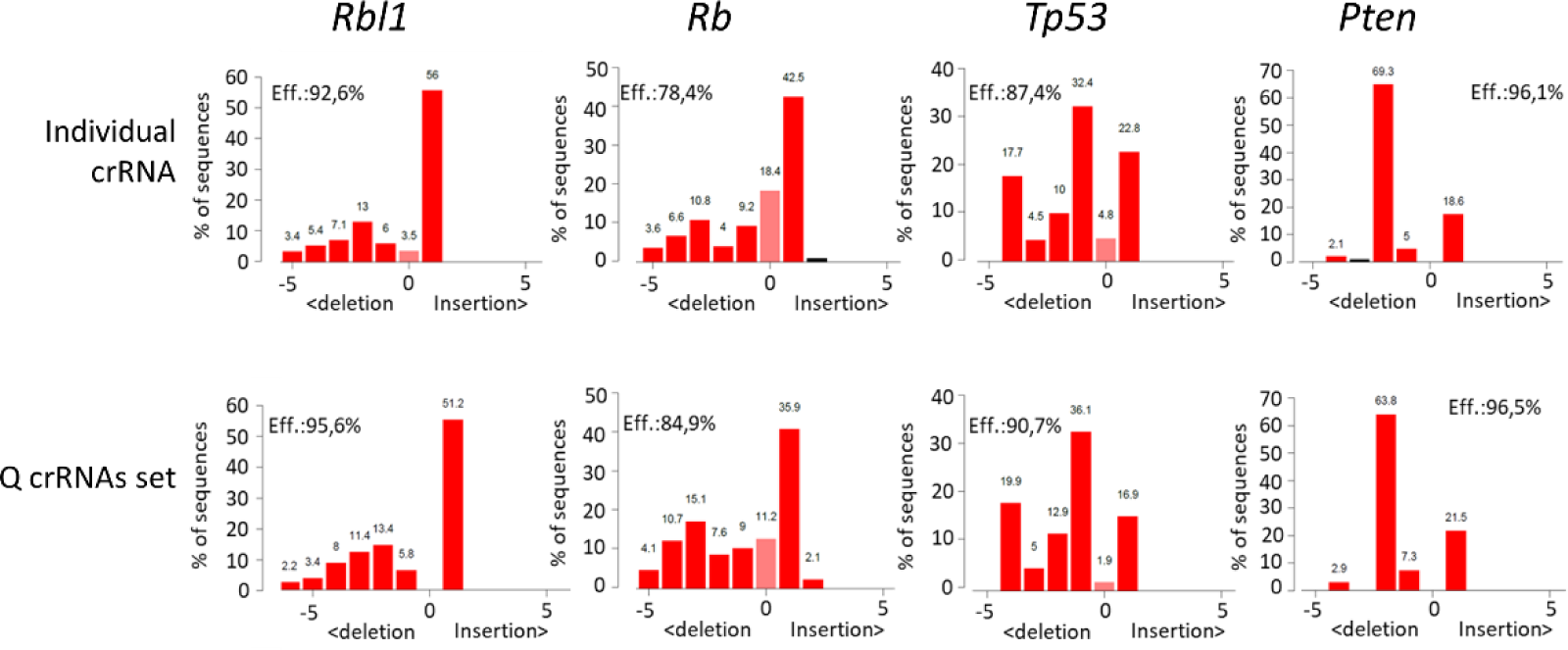
Efficient gene editing of the tumor suppressor genes Rbl1, Rb1, pTen, and Trp53 in Mouse Embryonic Fibroblasts (MEFs) Tracking of indels by decomposition (TIDE) analysis showing high editing efficiency of the four different gRNAs in RC::RFLTG MEFs. TIDE profiles of edited samples: the indel spectrum within ±10 bp from predicted cleavage site.

**Fig. S4.**
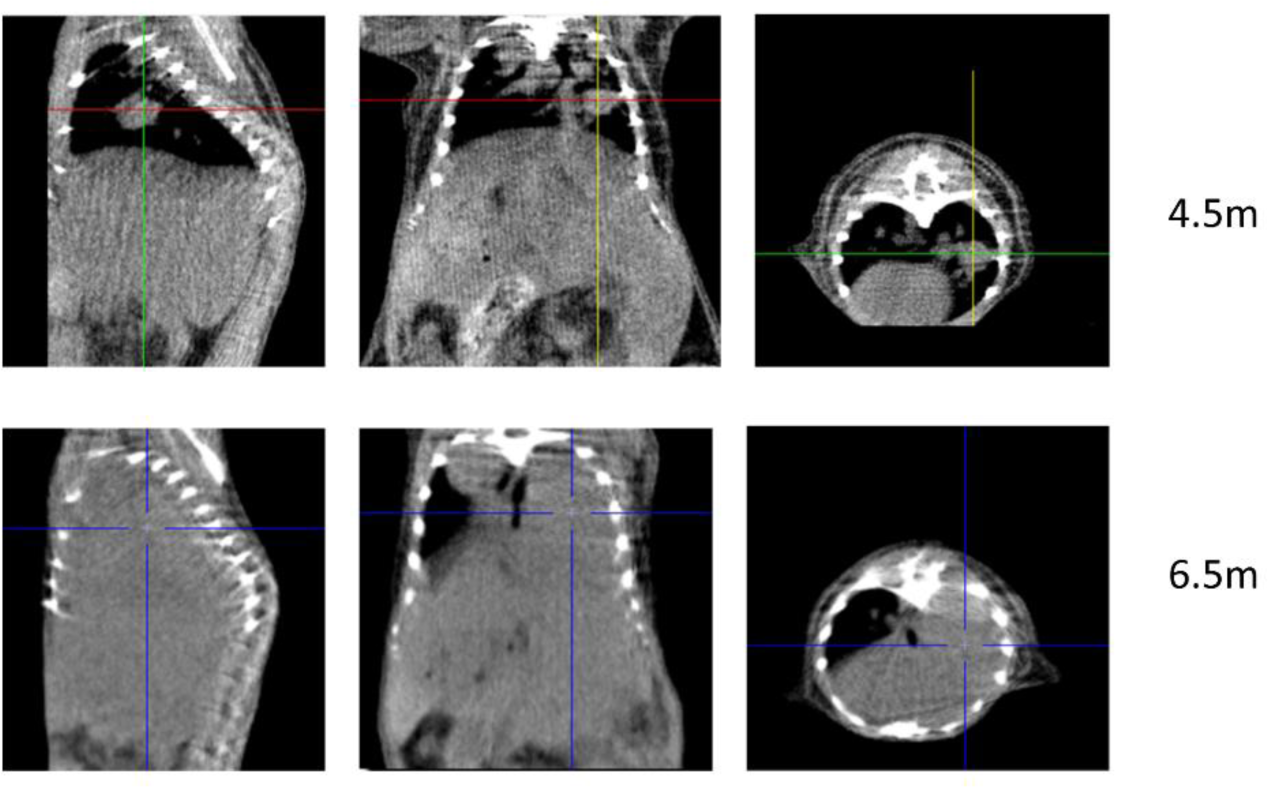
Monitoring lung tumor growth *in vivo*. Computed Tomography (CT) imaging of a RC::FLTG mouse treated with polyplexes (Qfrt Y4 Linear). Sagittal (left), coronal (center) and transverse (right) view images where a tumor is detected in the right lung growing over time (upper panels, CT scan taken 4.5 months after treatment, lower panels, 2 months later).

**Fig. S5.**
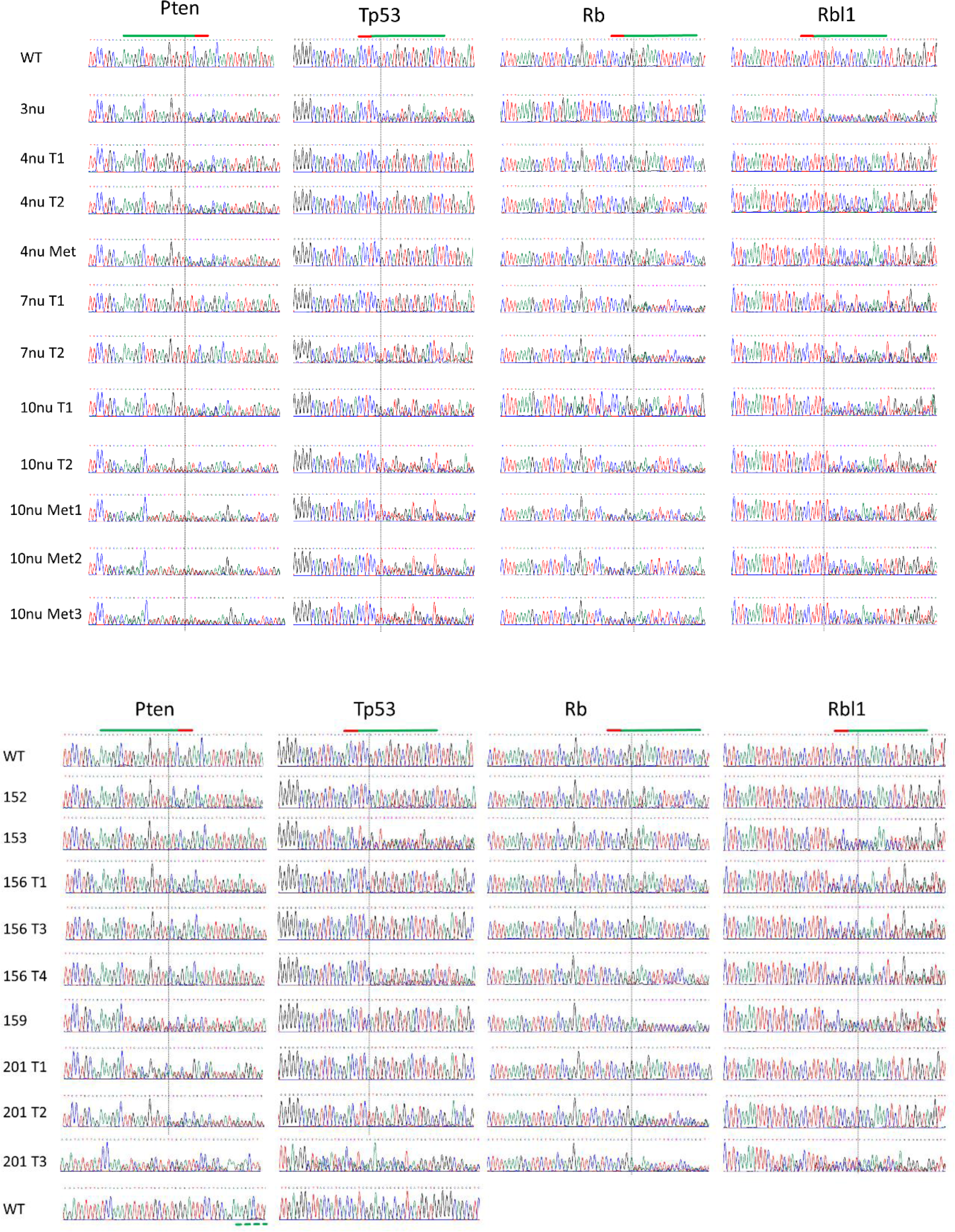
Tumor genotyping by Sanger Sequencing. Sequence chromatograms of the PCR amplified regions spanning the cleavage site for each tumor suppressor gene and each genotyped tumor. Tumors are identified by the ID# of the mouse. Immunodeficient mice are identified as “#nu” and metastases from these mice as “#nu Met#”. Where multiple tumors were genotyped from a single mouse, the tumor number is given as T#. The window shown for the *Pten* and *Trp53* gene sequences corresponding to the 201 T3 tumor has been adjusted to the larger size of the deletions. Sequences are shown paired with the corresponding control wild type sequences. Green line: gRNA target sequence; red line: PAM sequence. Dashed vertical line: predicted cut site for each guide. WT: control wild type sequence.

**Fig. S6.**
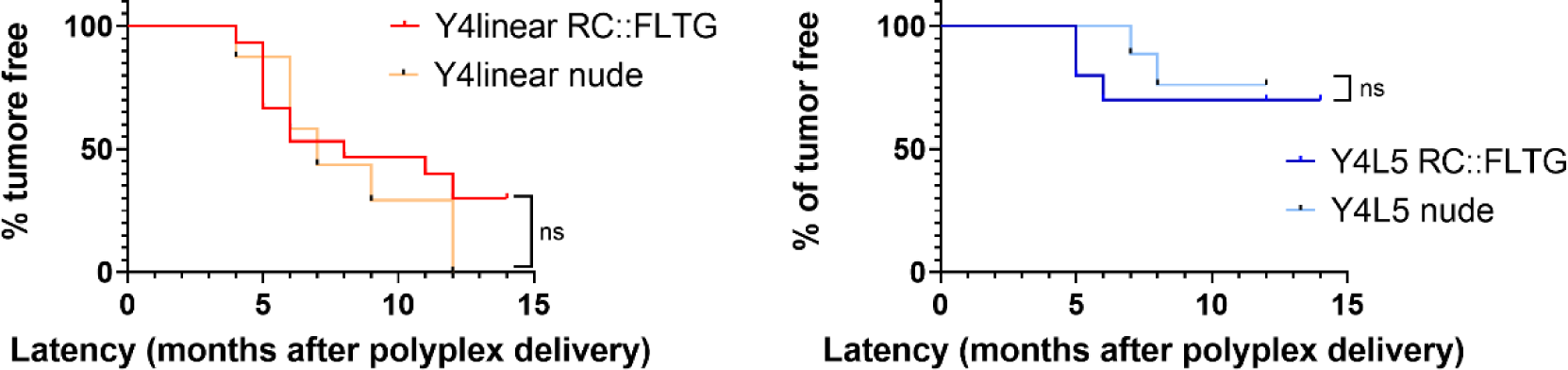
Survival curves comparison of polyplexes treated mice in immunocopetent and immunodeficient backgrounds. Kaplan-Meier survival curves of inmunodeficient (nude) or immunocompetent (RC::FLTG) mice treated with the indicated polymers (Y4 Linear (left) Y4 L5 (right). Differences in the survival curves are statistically non-significant (ns), determined by log-rank test.

**Fig. S7.**
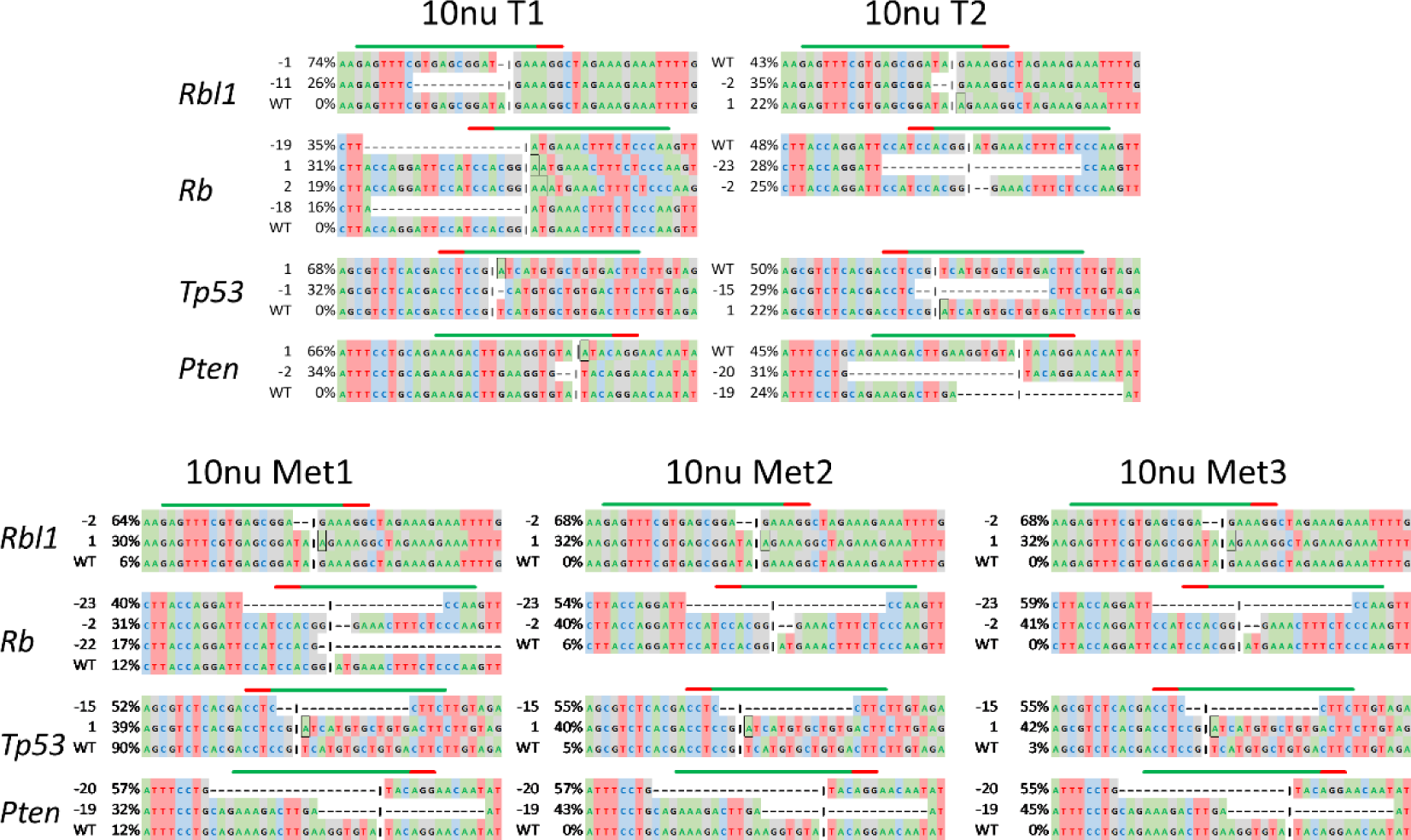
Primary lung tumor and liver metastasis genotyping by Sanger Sequencing. Sanger sequences of PCR amplicons of the four tumor suppressor genes corresponding to two different primary tumors (upper) from mouse 10nu and metastases (lower). All three metastases carry sets of indels that correspond to those found in a primary tumor (T2). T: tumor. Met: metastasis. Green line: gRNA target sequence; Red line: PAM sequence. Indel length and percentage shown on the left. Black rectangle boxes depict insertions. Cleavage site indicated by black vertical lines.

**Fig. S8.**
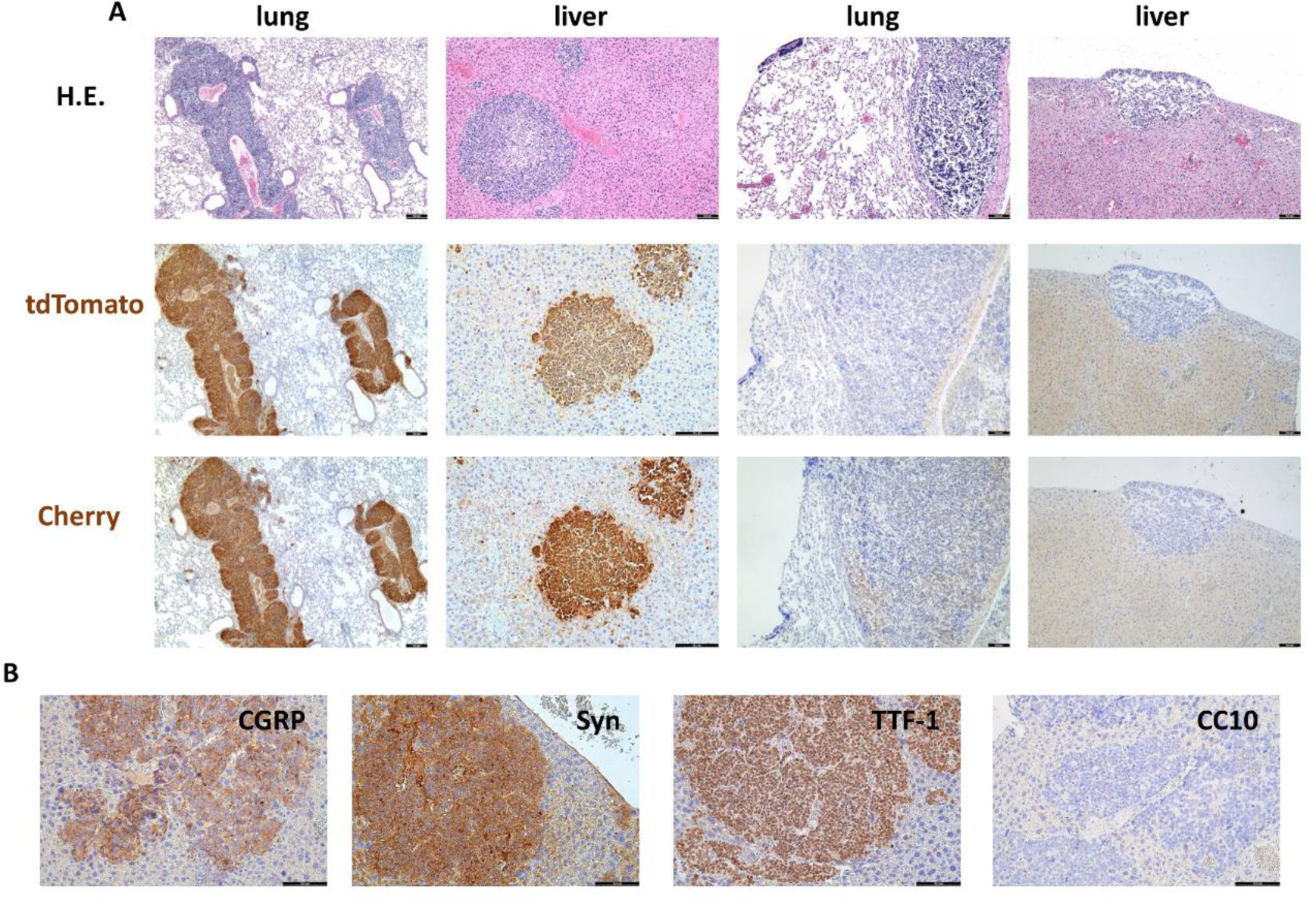
Characterization of SCLC mouse tumors and metastasis in RC::FLTG mice. (A) Immunohistochemical detection of tomato protein using tdTomato or mCherry antibodies in primary lung tumors and the corresponding liver metastases as quoted. B) Representative images of immunohistochemical staining of liver metastases for the quoted proteins: CGRP, Calcitonin Gene Related Protein; Syn, synaptphysin; TTF-1, Thyroid Transcription Factor-1; CC10, Clara cell secretory protein. Bars = 100 μm

## SUPPLEMENTARY TABLE

**Table S1.**
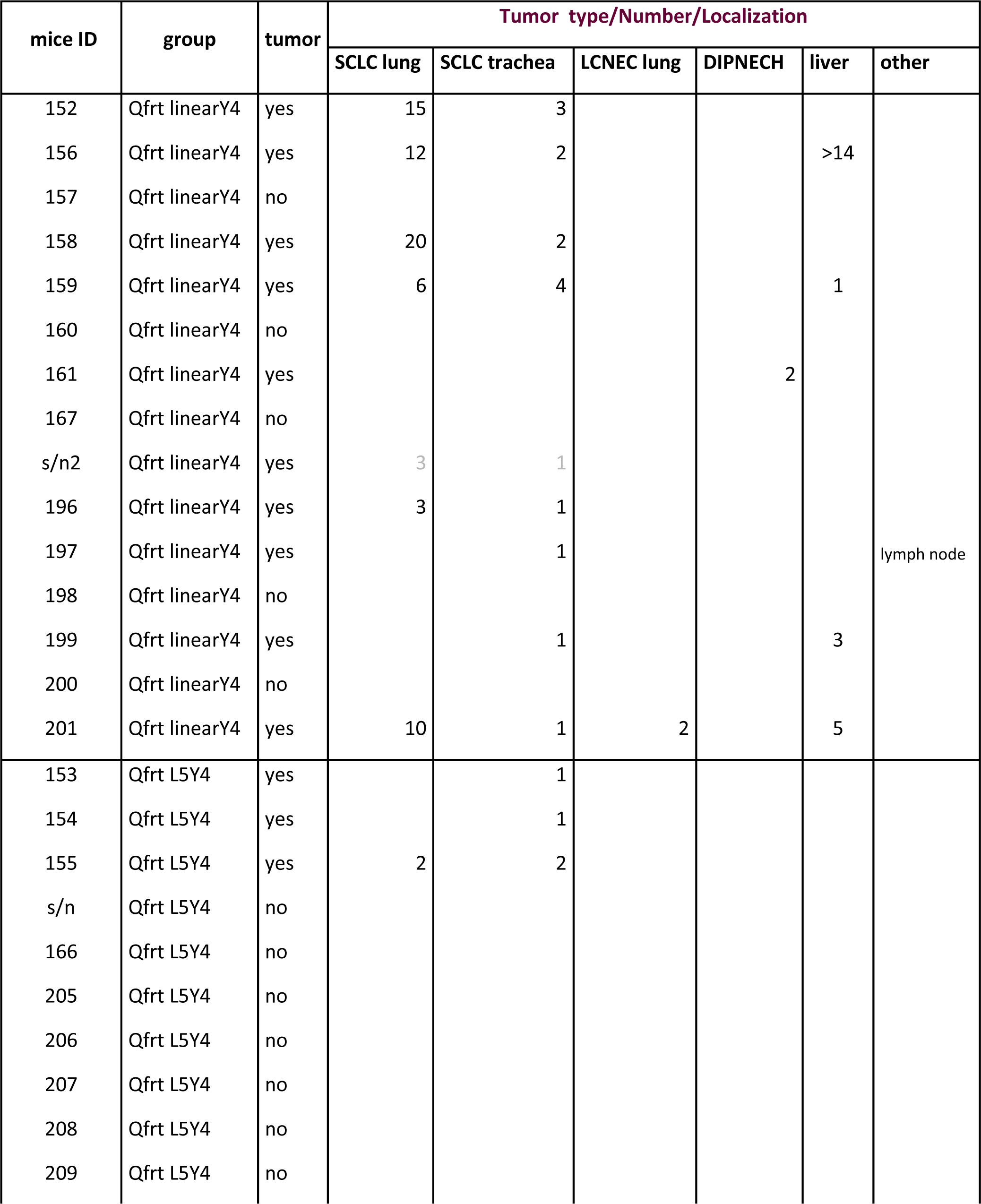
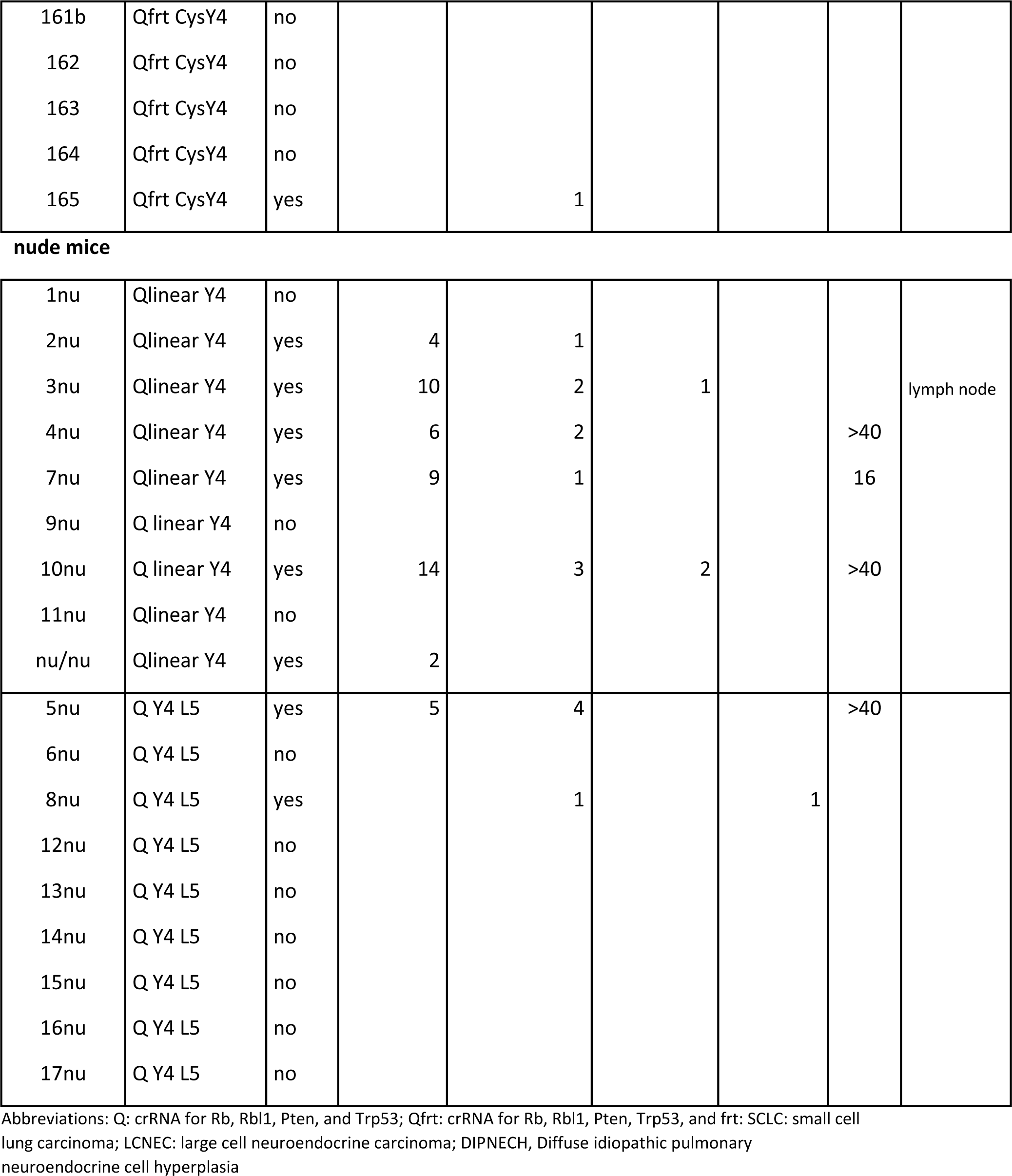
Tumors and Metastasis in Mice treated with ribolyplexes RC::FLTG mice.

| mice ID | group | tumor | Tumor type/Number/Localization |  |  |  |  |  |
| --- | --- | --- | --- | --- | --- | --- | --- | --- |
|  |  |  | SCLC lung | SCLC trachea | LCNEC lung | DIPNECH | liver | other |
| 152 | Qfrt linearY4 | yes | 15 | 3 |  |  |  |  |
| 156 | Qfrt linearY4 | yes | 12 | 2 |  |  | >14 |  |
| 157 | Qfrt linearY4 | no |  |  |  |  |  |  |
| 158 | Qfrt linearY4 | yes | 20 | 2 |  |  |  |  |
| 159 | Qfrt linearY4 | yes | 6 | 4 |  |  | 1 |  |
| 160 | Qfrt linearY4 | no |  |  |  |  |  |  |
| 161 | Qfrt linearY4 | yes |  |  |  | 2 |  |  |
| 167 | Qfrt linearY4 | no |  |  |  |  |  |  |
| s/n2 | Qfrt linearY4 | yes | 3 | 1 |  |  |  |  |
| 196 | Qfrt linearY4 | yes | 3 | 1 |  |  |  |  |
| 197 | Qfrt linearY4 | yes |  | 1 |  |  |  | lymph node |
| 198 | Qfrt linearY4 | no |  |  |  |  |  |  |
| 199 | Qfrt linearY4 | yes |  | 1 |  |  | 3 |  |
| 200 | Qfrt linearY4 | no |  |  |  |  |  |  |
| 201 | Qfrt linearY4 | yes | 10 | 1 | 2 |  | 5 |  |
| 153 | Qfrt L5Y4 | yes |  | 1 |  |  |  |  |
| 154 | Qfrt L5Y4 | yes |  | 1 |  |  |  |  |
| 155 | Qfrt L5Y4 | yes | 2 | 2 |  |  |  |  |
| s/n | Qfrt L5Y4 | no |  |  |  |  |  |  |
| 166 | Qfrt L5Y4 | no |  |  |  |  |  |  |
| 205 | Qfrt L5Y4 | no |  |  |  |  |  |  |
| 206 | Qfrt L5Y4 | no |  |  |  |  |  |  |
| 207 | Qfrt L5Y4 | no |  |  |  |  |  |  |
| 208 | Qfrt L5Y4 | no |  |  |  |  |  |  |
| 209 | Qfrt L5Y4 | no |  |  |  |  |  |  |
| 161b | Qfrt CysY4 | no |  |  |  |  |  |  |
| 162 | Qfrt CysY4 | no |  |  |  |  |  |  |
| 163 | Qfrt CysY4 | no |  |  |  |  |  |  |
| 164 | Qfrt CysY4 | no |  |  |  |  |  |  |
| 165 | Qfrt CysY4 | yes |  | 1 |  |  |  |  |
| **nude mice** |  |  |  |  |  |  |  |  |
| 1nu | Qlinear Y4 | no |  |  |  |  |  |  |
| 2nu | Qlinear Y4 | yes | 4 | 1 |  |  |  |  |
| 3nu | Qlinear Y4 | yes | 10 | 2 | 1 |  |  | lymph node |
| 4nu | Qlinear Y4 | yes | 6 | 2 |  |  | >40 |  |
| 7nu | Qlinear Y4 | yes | 9 | 1 |  |  | 16 |  |
| 9nu | Q linear Y4 | no |  |  |  |  |  |  |
| 10nu | Q linear Y4 | yes | 14 | 3 | 2 |  | >40 |  |
| 11nu | Qlinear Y4 | no |  |  |  |  |  |  |
| nu/nu | Qlinear Y4 | yes | 2 |  |  |  |  |  |
| 5nu | Q Y4 L5 | yes | 5 | 4 |  |  | >40 |  |
| 6nu | Q Y4 L5 | no |  |  |  |  |  |  |
| 8nu | Q Y4 L5 | yes |  | 1 |  | 1 |  |  |
| 12nu | Q Y4 L5 | no |  |  |  |  |  |  |
| 13nu | Q Y4 L5 | no |  |  |  |  |  |  |
| 14nu | Q Y4 L5 | no |  |  |  |  |  |  |
| 15nu | Q Y4 L5 | no |  |  |  |  |  |  |
| 16nu | Q Y4 L5 | no |  |  |  |  |  |  |
| 17nu | Q Y4 L5 | no |  |  |  |  |  |  |

Abbreviations: Q: crRNA for Rb, Rbl1, Pten, and Trp53; Qfrt: crRNA for Rb, Rbl1, Pten, Trp53, and frt; SCLC: small cell lung carcinoma; LCNEC: large cell neuroendocrine carcinoma; DIPNECH, Diffuse idiopathic pulmonary neuroendocrine cell hyperplasia

